# A sequence-based global map of regulatory activity for deciphering human genetics

**DOI:** 10.1101/2021.07.29.454384

**Authors:** Kathleen M. Chen, Aaron K. Wong, Olga G. Troyanskaya, Jian Zhou

## Abstract

Sequence is at the basis of how the genome shapes chromatin organization, regulates gene expression, and impacts traits and diseases. Epigenomic profiling efforts have enabled large-scale identification of regulatory elements, yet we still lack a sequence-based map to systematically identify regulatory activities from any sequence, which is necessary for predicting the effects of any variant on these activities. We address this challenge with Sei, a new framework for integrating human genetics data with sequence information to discover the regulatory basis of traits and diseases. Our framework systematically learns a vocabulary for the regulatory activities of sequences, which we call sequence classes, using a new deep learning model that predicts a compendium of 21,907 chromatin profiles across >1,300 cell lines and tissues, the most comprehensive to-date. Sequence classes allow for a global view of sequence and variant effects by quantifying diverse regulatory activities, such as loss or gain of cell-type-specific enhancer function. We show that sequence class predictions are supported by experimental data, including tissue-specific gene expression, expression QTLs, and evolutionary constraints based on population allele frequencies. Finally, we applied our framework to human genetics data. Sequence classes uniquely provide a non-overlapping partitioning of GWAS heritability by tissue-specific regulatory activity categories, which we use to characterize the regulatory architecture of 47 traits and diseases from UK Biobank. Furthermore, the predicted loss or gain of sequence class activities suggest specific mechanistic hypotheses for individual regulatory pathogenic mutations. We provide this framework as a resource to further elucidate the sequence basis of human health and disease.

## Introduction

Deciphering how regulatory functions are encoded in genomic sequences is a major challenge in understanding how genome variation links to phenotypic traits. Cell-type-specific regulatory activities encoded in elements such as promoters, enhancers, and chromatin insulators are critical to defining the complex expression programs essential for multicellular organisms, like those affecting cell lineage specificity and development. The majority of disease-associated variants from genome-wide association studies (GWAS) are located in noncoding regions^1^ and may perturb regulatory elements, yet without knowing how changes in sequence affect regulatory activities we cannot predict the impact of these variants and uncover the regulatory mechanisms contributing to complex diseases and traits. Different variants in the same region can have distinct regulatory consequences and resulting phenotypic effects, as shown by mutations in enhancer regions of SHH^2^: for instance, a variant may turn off the expression of a gene critical for early development in specific tissue and location, while other variants in the same region may increase enhancer activity or have no effect at all.

Substantial progress has been made in the experimental profiling and integrative analysis of epigenomic marks, such as histone marks and DNA accessibility, across a wide range of tissues and cell types^3–5^. Histone marks are commonly used to identify regulatory elements; for example, H3K4me3 can indicate active promoter regions and H3K27ac/H3K4me1 can indicate active enhancer regions. Moreover, histone marks and chromatin accessibility can be integrated with chromatin state models^6–10^. These works have been instrumental to annotating the genome with regulatory elements across many tissues.

At the same time, deep learning sequence modeling techniques have been successfully applied to learn sequence features that are predictive of transcription factor binding and histone modifications^11–17^. These models are powerful tools for inferring the impact of sequence variation at the chromatin level. However, each chromatin-level prediction can only inform a very specific aspect of sequence--for example, whether a variant causes an increase or decrease of C/EBP-β binding. We continue to lack a global, integrative view of sequence regulatory activities, including all major aspects of cis-regulatory functions, such as tissue-specific or broad enhancer and promoter activities. This limits our ability to interpret the integrated effects of all chromatin-level perturbations caused by genomic variants and determine their impact on human health and diseases.

We address this challenge by creating a global map for sequence regulatory activity based on a new deep-learning-based framework called Sei. This framework introduces a new sequence model that predicts a comprehensive compendium of 21,907 publicly available chromatin profiles--the broadest set to-date--and uses the model to quantitatively characterize regulatory activities for any sequence with a novel vocabulary of sequence classes. Sequence classes cover diverse types of regulatory activities, such as promoter or cell-type-specific enhancer activity, across the whole genome by integrating sequence-based predictions from histone marks, transcription factors, and chromatin accessibility across a wide range of cell types. For example, ‘embryonic stem cell-specific enhancer’ sequence class activity may be estimated from the predicted binding of multiple transcription factors including Pou5F1, Sox2, and Nanog, as well as various histone marks, on a sequence. Importantly, sequence classes can be used to both classify and quantify the regulatory activities of any sequence based on predictions made by the deep learning sequence model. Therefore, sequence classes allow for the quantitative mapping of any mutation to its impact on cell-type-specific regulatory activities.

The Sei framework thus provides an interpretable and systematic integration of sequence-based regulatory activity predictions (intrinsic information, based on sequence function) with human genetics data (extrinsic information, based on variant-phenotype association) for discovering the regulatory basis of human traits and disease. We applied our framework to characterize disease- and trait-associated regulatory disruptions by combining sequence class information and UK Biobank GWAS data. Sequence classes provide a non-overlapping partitioning of heritability in GWAS by regulatory activity, which we use to profile the regulatory architecture of 47 diseases and traits in UK Biobank GWAS^18^.

Moreover, variant effect prediction at the sequence-class-level newly enables the interpretation of regulatory mechanisms for individual disease mutations and can differentiate between gain-of-function and loss-of-function regulatory mutations. The regulatory and tissue-specific view provided by sequence classes suggests potential new mechanisms for individual disease-associated variants: for example, we used sequence classes to link mutations in blood-related diseases with previously unknown mechanisms to the malfunctioning of cell-type-specific enhancers.

We provide the Sei framework as a resource for systematically classifying and scoring any sequence and variant with sequence classes, additionally providing the Sei model predictions for the 21,907 chromatin profiles underlying the sequence classes. The framework can be run using the code at https://github.com/FunctionLab/sei-framework, and a user-friendly web server is available at hb.flatironinstitute.org/sei.

## Results

### Developing a comprehensive sequence model for 21,907 chromatin profiles

To capture the widest range of sequence features that are predictive of regulatory activities, we first developed a new deep learning sequence model, which we refer to as the Sei model, that enables the base-level interpretation of sequences by predicting 21,907 genome-wide cis-regulatory targets--including peak calls from 9,471 transcription factor profiles, 10,064 histone mark profiles and 2,372 chromatin accessibility profiles--with single nucleotide sensitivity. The majority of this data (19,905 profiles) is from the Cistrome Project^5^, a resource that uniformly processes and annotates public ChIP-, DNase-, and ATAC-seq datasets, and the remaining chromatin profiles were processed by the ENCODE^3^ and Roadmap Epigenomics^4^ projects. The Sei model encompasses an estimated ∼1000 non-histone DNA-binding proteins (which we refer to as transcription factors), 77 histone marks, and chromatin accessibility across >1300 cell lines and tissues (Supplementary Files 1, 2).

To efficiently predict 21,907 chromatin profiles from sequence, we designed a novel model architecture (Supplementary Figure 1) and improved our training pipeline. The Sei model uses a new residual-block architecture with a dual linear and nonlinear path design: the linear path allows for fast and statistically efficient training, while the nonlinear path offers strong representation power and the capability to learn complex interactions. For scaling and performance, we introduced a layer of spatial basis functions, which integrates information across spatial locations with much higher memory efficiency than fully connected layers. The model takes as input a 4kb length sequence and predicts the probabilities of 21,907 targets at the center position. The model is trained on chromatin profile peak calls, which are binary (presence/absence), but the model output is continuous, representing probabilities of peaks. Our model training pipeline was updated to improve training speed and performance by using on-the-fly sampling, which reduces overfitting by generating new training samples for every training step.

The model achieved an average area under the receiver-operating characteristic (AUROC) of 0.972 and average area under the precision-recall curve (AUPRC) of 0.409 across all 21,907 chromatin profiles (Supplementary Figure 2). In addition to accurately predicting individual profiles, the predictions also recapitulated the correlation structure of these profiles, which indicates that the Sei model is able to capture the co-localization patterns of chromatin profiles (Supplementary Figure 3). Furthermore, the Sei model also improved over our best previously published model, DeepSEA “Beluga”^13^, on the 2002 chromatin profiles predicted by both models by 19% on average (as measured by AUROC/1-AUROC, Supplementary Figure 4).

Therefore, the Sei model is the most comprehensive chromatin-level sequence model to-date, and offers an expansive new resource for sequence and variant interpretation.

### Defining sequence classes using a sequence model from whole genome sequences

Next, we applied the Sei model to develop a global, quantitative map from genomic sequences to specific classes of regulatory activities, which we term sequence classes, by integrating the wide range of chromatin profiles predicted by Sei. Sequence classes are therefore mapped directly from sequence, and each sequence class represents a distinct program of regulatory activities across tissues and cell types as covered by the Sei model. Furthermore, sequence classes allow for the mapping of any sequence to quantitative scores that represent a broad spectrum of regulatory activities.

To cover the whole spectrum of sequence activities, we identified sequence classes from Sei predictions for 30 million sequences uniformly tiling the whole genome (4kb windows with 100bp step size). We visualized the global structure of sequence regulatory signals as represented by the model’s chromatin profile predictions with nonlinear dimensionality reduction techniques^19, 20^ (Figure 1) and applied Louvain community clustering^21^ to these predictions to categorize the 30 million sequences into 40 sequence classes (Figure 1a).

**Figure 1.**
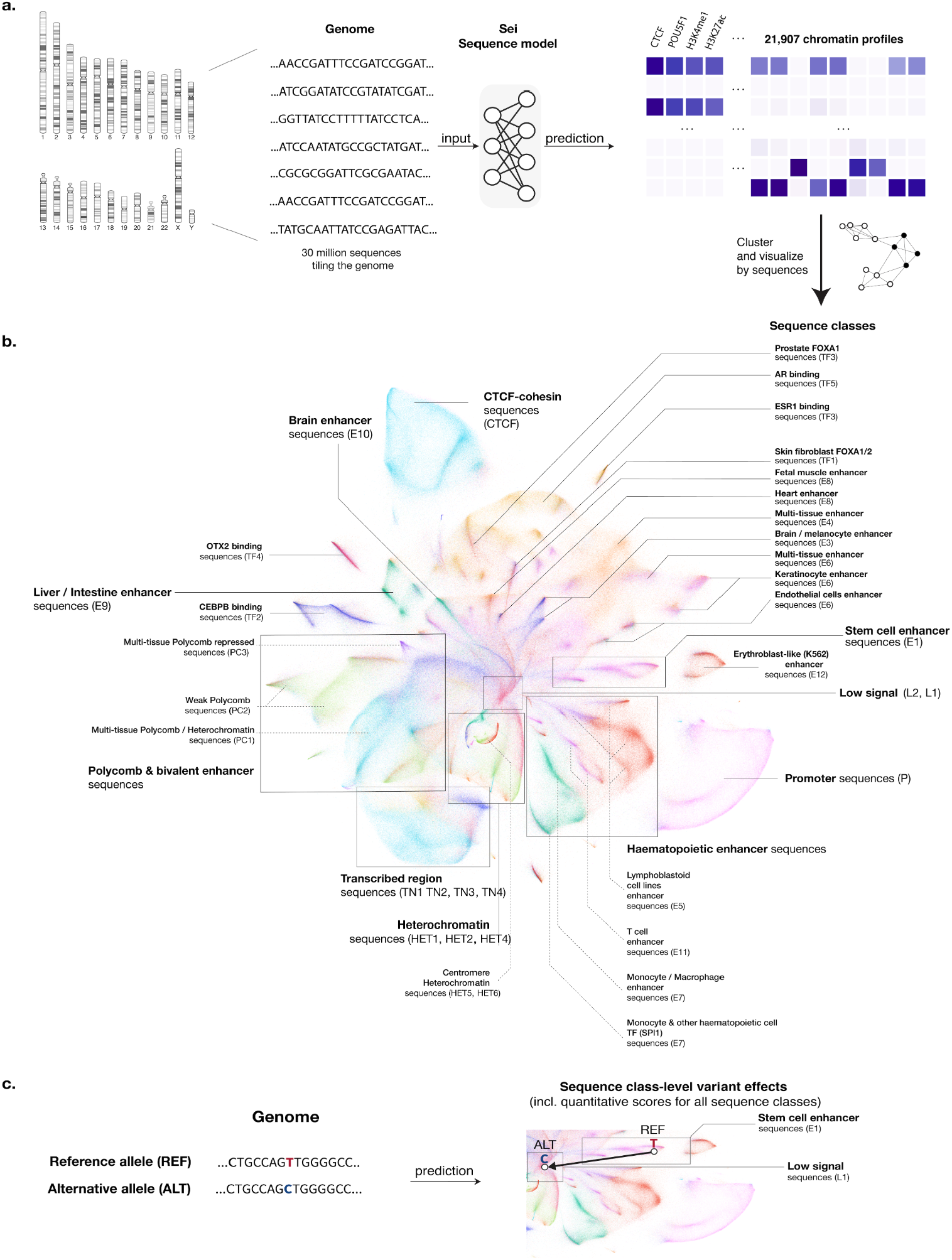
Mapping the global regulatory landscape of genomic sequences. **a,** Overview of the Sei framework for systematic prediction of sequence regulatory activities. Sequence classes are extracted from the predicted chromatin profiles of 30 million sequences evenly tiling the genome. The predictions were made by Sei, a new deep convolutional network sequence model trained on 21,907 chromatin profiles. Specifically, classes are identified by applying Louvain community detection to the nearest-neighbor graph of 180 principal components extracted from the predictions data. **b,** Visualizing the global regulatory landscape of human genome sequences discovered by this approach with UMAP. Major sequence classes include cell-type-specific enhancer classes, CTCF-cohesin, promoter, TF-specific, and heterochromatin/centromere classes. **c,** This framework is further applied to predict sequence-class-level genome variant effects, quantified by changes in sequence class scores.

This visualization of human genome sequences demonstrates the global organization of sequence regulatory activities (Figure 1b). The center of the visualization contains sequences with weak or no regulatory activity based on histone mark and TF enrichment, and sequences with specific regulatory activities radiate outwards, establishing a continuum from no activity to strong specific activity. Different branches of sequences are enriched in distinct chromatin modifications and transcription factors, and sequences with similar regulatory activities are grouped together. For example, tissue-specific enhancer sequences were predominantly grouped by tissue in the visualization (Figure 1b). In addition, sequences with repressive Polycomb marks were spatially adjacent to H3K9me3-marked heterochromatin sequences (Figure 1b), reflecting their extensive crosstalk in epigenetic silencing^22–24^. Notably, promoter-proximal and CTCF-cohesin binding sequences form two well-defined clusters that are separated from other sequences, which may reflect the distinct nature of these activities (Figure 1b).

The sequence classes identified from whole genome sequences recapitulate the sequence organization shown in the visualization and provide a basis for summarizing sequence activities globally and are robust to clustering parameter choices (Supplementary Figures 5, 6). To facilitate intuitive interpretation of sequence classes, we named them based on the corresponding enrichment of cis-regulatory profiles (Figure 2a, Supplementary Figures 7-12, Supplementary File 3); specifically, we label each sequence class with a functional group acronym and index denoting the rank of the sequence class within the group (Supplementary Figure 13, e.g. E1 encompasses a larger proportion of the genome than E2). Because genomic sequences encode their regulatory activity programs across all cell types, sequence classes also show distinct activity patterns across cell types and tissues. We label sequence classes primarily based on their active, cell-type-specific regulatory activities--in particular, promoter and enhancer activities. Therefore, sequence classes that are not labeled as enhancer (‘E’) or promoter (‘P’) generally lack enhancer or promoter activity in any cell type predicted by Sei.

**Figure 2.**
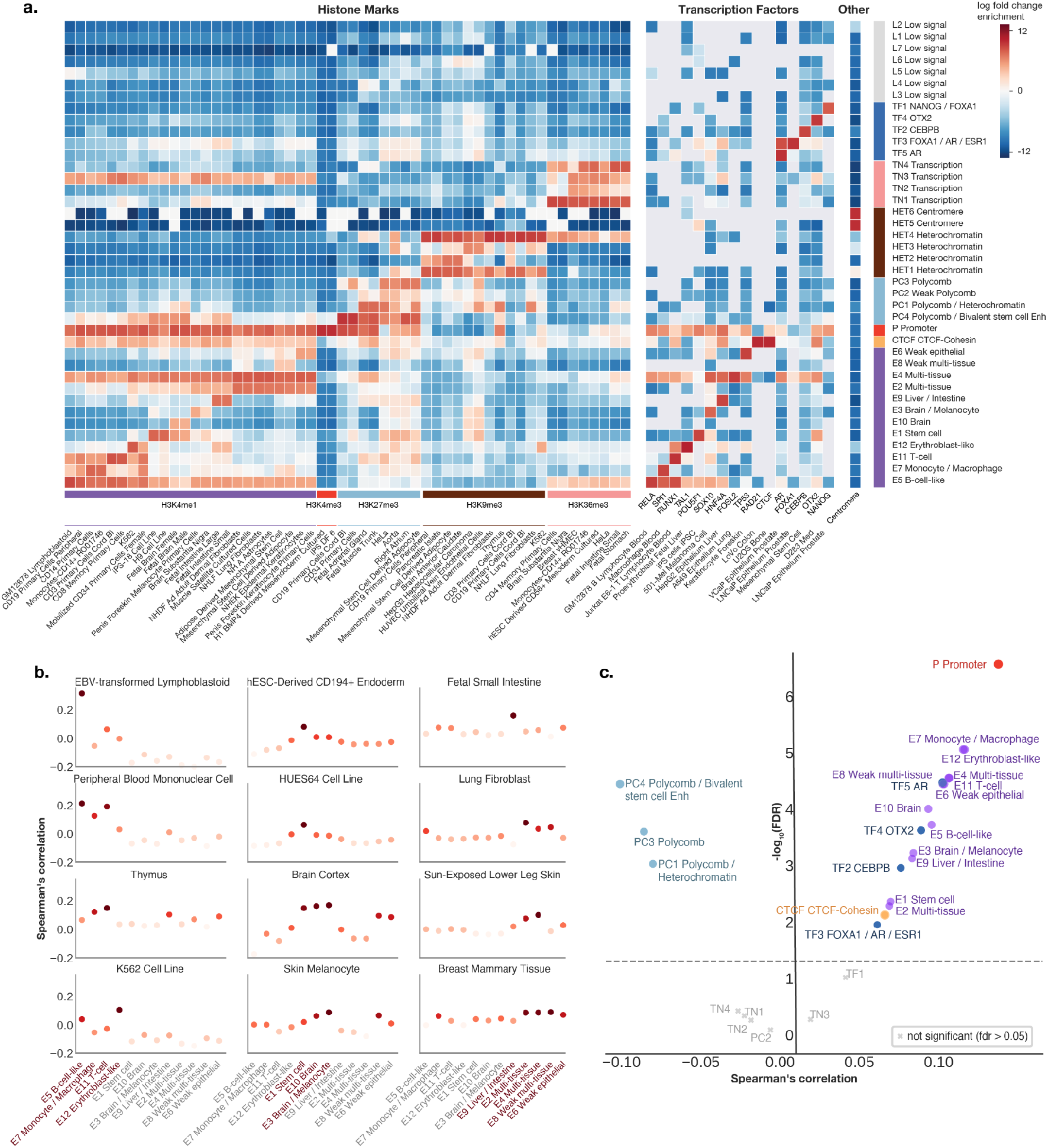
Sequence classes predict cell-type-specific regulatory activities and directional, expression-altering variant effects. **a,** Sequence-class-specific enrichment of histone marks, transcription factors, and repeat annotations. Log fold change enrichment over genome-average background is shown in the heatmap. No overlap is indicated by the gray color in the heatmap. Top 1-2 histone mark and TF annotation enrichments were selected for each sequence class. **b,** Enhancer sequence classes near transcription start sites are correlated with cell-type-specific gene expression in the applicable tissue or cell types (see Methods). The y-axis shows the Spearman correlation between the proportion of each sequence class annotation within 10kb of TSS and the tissue-specific differential gene expression (fold over tissue-average). **c,** Regulatory sequence-class-level variant effects are predictive of directional GTEx variant gene expression effects. The x-axis shows Spearman correlations between the predicted sequence-class-level variant effects and the signed GTEx variant effect sizes (slopes) for variants with strong predicted effects near transcription start sites (Methods) and the y-axis shows the corresponding log10 p-values. All colored dots are above the Benjamini-Hochberg FDR < 0.05 threshold.

**Figure 3.**
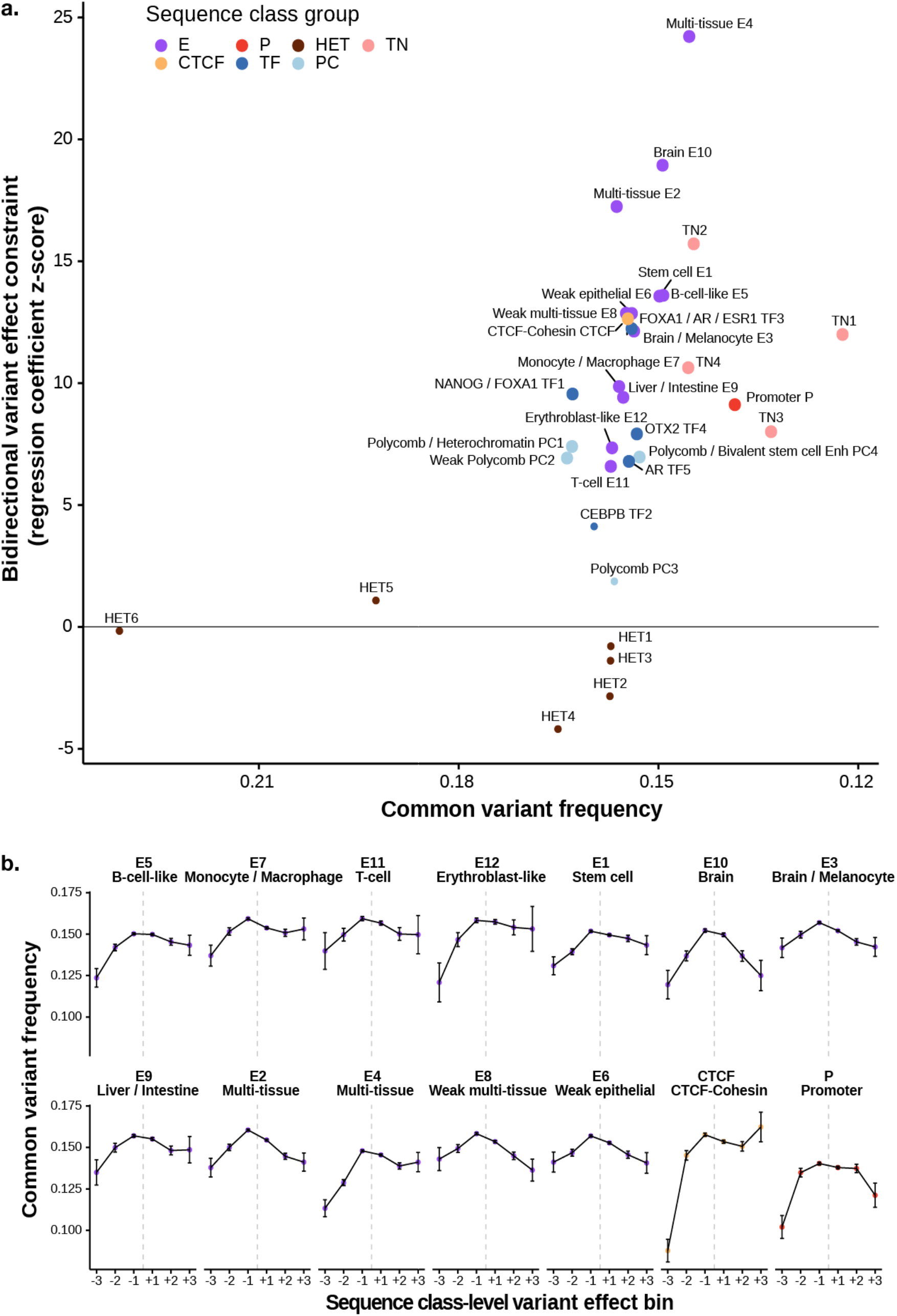
Variants with strong regulatory sequence class effects show negative selection signatures. **a,** Scatter plot for allele-frequency-based analysis of each sequence class. The x-axis shows 1 - common variant frequency (allele frequency > 0.01) across all 1000 Genome variants per sequence class, and the y-axis shows the bidirectional variant effect constraint z-score, which is computed based on logistic regressions predicting common variant (allele frequency > 0.01) from sequence-class-level variant effect score for both positive and negative effects (Methods). Sequence classes with significant (Bonferroni-Hochberg FDR<0.05) bidirectional variant effect constraint are indicated with larger dots. ‘L’ sequence classes are excluded due to lack of interpretation for their sequence-class-level variant effect scores. **b,** Comparison of common variant frequencies for 1000 Genomes variants assigned to different sequence classes and variant effect bins. The common variant threshold is >0.01 allele frequency across the 1000 Genomes population. Error bars show +/- 1 standard error (SE). The sequence-class-level variant effects are assigned to 6 bins (+3: top 1% positive, +2: top 1%-10% positive, +1: top 10% -100% positive, -1: top 10% -100% negative, -2: top 1%-10% negative, -3: top 1% negative).

In summary, sequence classes contain 1 ‘P’ promoter class, which is most strongly enriched in the active promoter histone mark H3K4me3 across all cell types (Figure 2a, Supplementary Figure 7); 12 ‘E’ enhancer classes, which are strongly enriched in enhancer histone marks, such as H3K4me1 and H3K27ac, and transcription factors relevant to their activities in select cell types (e.g. PU.1/Spi1 in the E7 monocyte/macrophage enhancer class, HNF4-α in E9 liver/intestine, and Sox2/Nanog/Pou5f1 in E1 stem cell), and often display repressive H3K27me3 marks in inactive cell types (Figure 2a, Supplementary Figures 8-10, Supplementary File 3); 1 ‘CTCF’ sequence class, which is strongly enriched in CTCF and cohesin (Figure 2a, Supplementary File 3); 5 ‘TF’ sequence classes, which are enriched in a few specific transcription factors (e.g. CEBPB sequence class) but have weak or no enhancer mark enrichment (Figure 2a, Supplementary File 3); 4 ‘PC’ Polycomb classes, which are enriched in the Polycomb-repressed region mark H3K27me3 and generally not enriched in active promoter or enhancer marks (Figure 2a, Supplementary Figure 10); 6 ‘HET’ heterochromatin classes, which are enriched in the heterochromatin mark H3K9me3 (Figure 2a, Supplementary Figure 11); 4 ‘TN’ sequence classes, which are enriched in transcription elongation marks H3K36me3 or H3K79me2 (Figure 2a, Supplementary Figure 12); and finally, 7 ‘L’ (low signal) sequence classes, which are not strongly enriched in any of the above marks (Figure 2a). As a whole, the 40 sequence classes cover >97.4% of the genome (Supplementary Figure 13).

Beyond classifying genomic sequences to sequence classes, we define sequence class scores to provide a global and quantitative representation of sequence regulatory activities. This for the first time allows us to (1) predict the regulatory activity for any sequence and (2) quantify the changes in regulatory activity caused by any sequence variant. Sequence class scores summarize predictions for all 21,907 chromatin profiles based on weights specific to each sequence class, which are computed by projecting Sei predictions onto unit-length vectors that point to the center of each sequence class. Sequences that score highly for a particular sequence class have high predictions for the chromatin profiles associated with that class. Sequence class scores thus allow for the quantification of the regulatory activity of any sequence, where the impact of a variant is represented by the difference between the sequence class scores for the reference and alternative alleles. Importantly, this capability is only allowed by modeling the sequence dependencies of sequence class activities and cannot be directly obtained from chromatin profiling data alone.

### Enhancer sequence classes predict tissue-specific gene expression

The group of sequences that are likely most impactful to tissue-specific gene expression regulation are the enhancer (‘E’) sequence classes, thus here we assessed the association of enhancer sequence class scores with tissue-specific gene expression.

In the visualization of sequence regulatory activities, sequence classes with different cell type- and tissue-specific enhancer activities are localized to distinct subregions (Figure 1b). ‘E’ sequence classes capture both specific and broad enhancer activities. Based on enhancer mark enrichment (Supplementary Figures 8, 9), E7 is specific for monocyte/macrophage, E11 is specific for T-cell, E5 is specific for lymphoblastoid/B-cell-like cell lines, E9 is specific for liver and intestine, E1 is specific for embryonic stem cells & induced pluripotent stem cells, and E10 and E3 are specific for brain (Figure 1, 2; all enrichments stated are significant with p<2.2e-16, Fisher’s exact test, two-sided). In contrast, broad enhancer sequence classes can either encompass enhancer activity in similar cell types across different tissues, such as fibroblast (E2) and epithelial (E6) cell types (Supplementary Figures 8, 9), or encompass enhancer activity in many different cell types; for example, E4 is enriched in fibroblast, muscle, astrocytes, osteoblast, epithelial, and other cell types. Sequence class enhancer activities are also supported by the enrichment of relevant chromatin states^3^ and DNase I hypersensitive sites^25^ across tissues and cell types (Supplementary Figures 14, 15). Consistent with their predicted enhancer activities, the coverage of ‘E’ sequence class annotations within a 10kb window to transcription start sites (TSS) are correlated with the differential expression patterns of these genes in the corresponding cell types over the tissue-average (Figure 2b).

Since sequence class scores allow us to systematically predict the effects of variants on higher-level regulatory functions, we can estimate whether a given variant diminishes, maintains, or increases the enhancer activity of a sequence based on the difference between the sequence class scores for the reference and alternative alleles. Evaluated on GTEx eQTL data^26^, we found that variants predicted to increase ‘E’ sequence class activity were significantly positively correlated with higher gene expression, whereas those predicted to increase ‘PC’ sequence class activity were significantly negatively correlated with gene expression--consistent with the expected repressive role of ‘PC’ sequence class activities (Figure 2c). Moreover, when only analyzing fine-mapped eQTLs^27^ with high posterior inclusion probability (>0.95), we observed higher correlations with overall comparable levels of significance (Supplementary Figure 16). Therefore, sequence classes can distinguish the effects of variants on gene expression based on their consequences in regulatory activities.

### Regulatory sequence classes are under evolutionary constraints

Variants that alter regulatory activities of sequences often disrupt gene regulation and are therefore expected to impact human health and disease. We tested this expectation by comparing human population genome variant allele frequencies^28^ based on the sequence class in which each variant is located and the predicted variant effect on that sequence class. Indeed, we found that variants localized in regulatory sequence classes (E-, P-, and CTCF-) have lower common variant frequency than variants in other sequence classes, and therefore showed higher overall negative selection constraint (Figure 3a, x-axis). More importantly, variants predicted to strongly perturb regulatory sequence classes had significantly lower common variant frequencies than variants that weakly perturb these classes (measured by bidirectional variant effect constraint, Figure 3a y-axis, see also Figure 3b, Methods). This is therefore consistent with the hypothesis that disruption of regulatory sequence class activities has a major negative impact on fitness, which we refer to as a negative selection signature.

Specifically, we observed strong negative selection signatures for variants assigned to all E, CTCF and P sequence classes (Figure 3). Multi-tissue enhancer sequence classes E4 and E2 and the brain enhancer sequence class E10 show the strongest association of predicted sequence-class-level variant effects and common variant frequencies. Notably, for the CTCF sequence class, only negative variant effects--decreasing sequence class activity--appear to be under very strong constraints, suggesting that CTCF sites are generally tolerant to positive effect mutations that further increase CTCF binding. This is in contrast to the generally deleterious impact of both increase and decrease of enhancer and promoter activities. As expected, TN sequence classes, which overlap with protein-coding regions, are among the sequence classes with the lowest allele frequency (Supplementary Figure 17).

In contrast, those assigned to HET, PC, TF, and L sequence classes generally did not show strong negative selection signatures and had higher overall common variant frequencies (Supplementary Figure 17). Importantly, this does not suggest that Polycomb or transcription factors are inessential: the HET, PC, TF, and L classes generally do not show strong enhancer or promoter histone mark enrichment in any cell type (with the exception of bivalent marks in stem cells observed in PC4), and thus they are expected to play less major roles in gene expression regulation. However, Polycomb-related regulation is likely critical for ‘E’ and ‘P’ sequence classes, which are often Polycomb-repressed in some cell types but enhancers or promoters in other cell types (Supplementary Figures 7-10). Similarly, we expect that TF binding plays a central role in ‘E’ classes that are highly enriched in relevant TFs (Figure 2a, Supplementary File 3).

Therefore, sequence classes show distinct evolutionary constraints, and ‘E’ enhancer sequence classes show the strongest bidirectional constraints. This suggests that both increases and decreases of enhancer activity are expected to lead to deleterious effects on fitness, highlighting the importance of precisely controlling gene expression.

### Sequence classes elucidate the tissue-specific regulatory architecture of GWAS traits

The population allele frequency analysis on sequence classes suggest that variants perturbing regulatory sequence class activities are likely involved in human health and disease. Therefore, to explore this hypothesis, we used GWAS data to delineate the genetic contribution of each sequence class to diseases and traits.

Partitioned heritability from LD score regression (LDSR) has been a powerful tool for understanding the genetic architecture of diseases and traits using GWAS summary statistics^29^, including identifying enrichment of disease heritability in regulatory elements ^29, 30^. Previous applications of LDSR use overlapping annotations,^29–31^ which allows for the joint analysis of heritability contribution across a wide range of annotations and has generated significant insight into a wide range of GWAS studies; however, such analyses cannot unambiguously partition heritability across annotations. Because sequence classes are both non-overlapping and cover nearly the entire genome, they provide a clear and more easily interpretable picture of the regulatory architecture of diseases and traits. To show this, we estimated the proportion of heritability explained by each sequence class for 47 GWAS traits in UK Biobank (UKBB)^18, 32^ (Methods). Specifically, we applied LDSR and used a conservative estimate of the proportion of heritability, subtracting one standard error and lower-bounding by 0. Our analysis of UKBB GWAS revealed genetic signatures of sequence-class-specific regulatory functions (Figure 4, Supplementary File 4).

**Figure 4.**
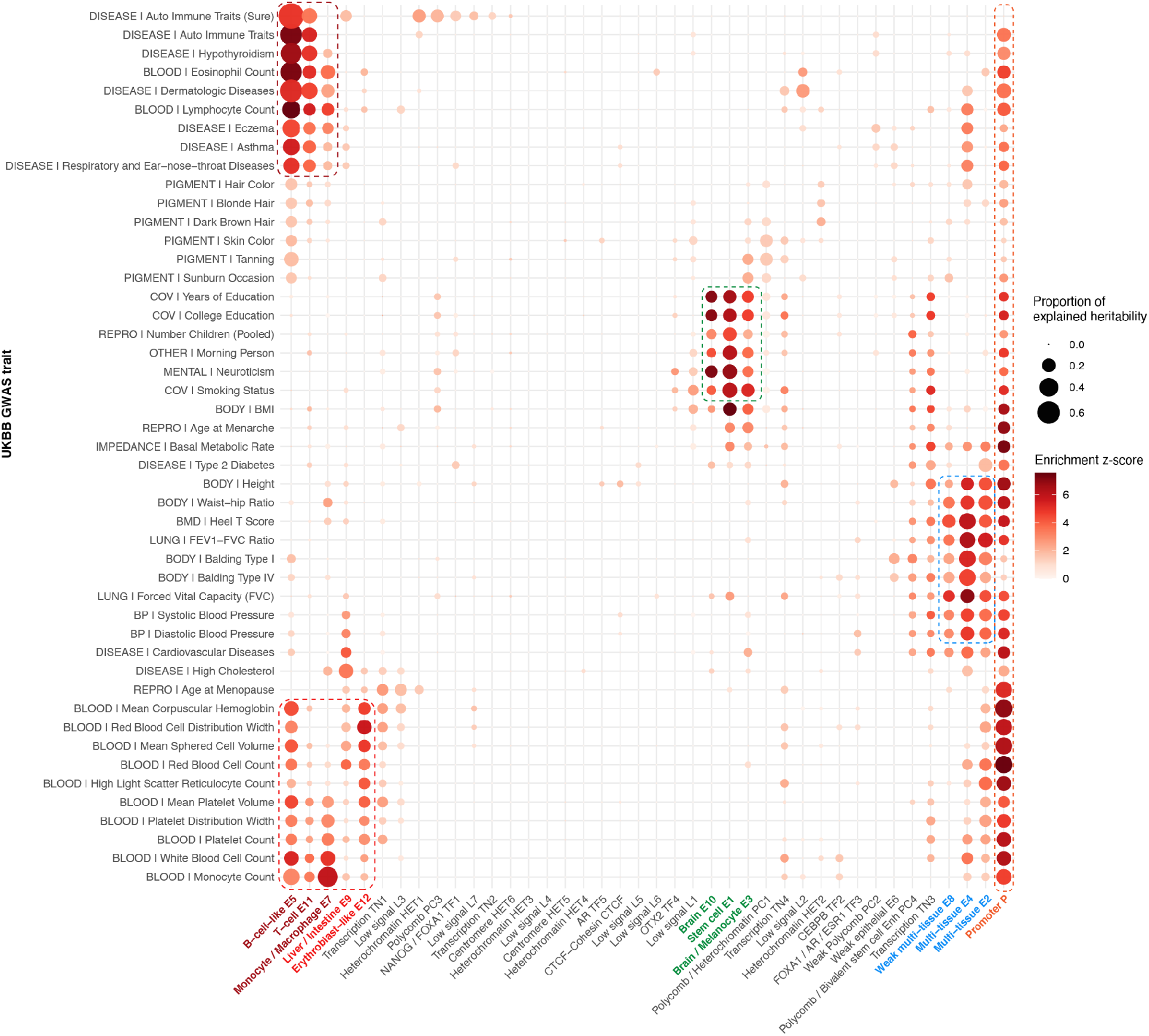
Sequence-class-based partitioning of GWAS heritability shows trait associations with tissue-specific regulation. Partitioned genome-wide heritability in UKBB GWAS with all 40 sequence classes. The size of the dot indicates the proportion of heritability estimated from LDSR, which is conservatively estimated as one standard error below the estimated heritability proportion (bounded by 0). The color of the dot indicates the significance z-score of the fold enrichment of heritability relative to the proportion of all SNPs assigned to the sequence class (bounded by 0). Colored boxes indicate traits associated with blood (red), brain (green), multiple tissues (blue) and promoters (orange).

**Figure 5.**
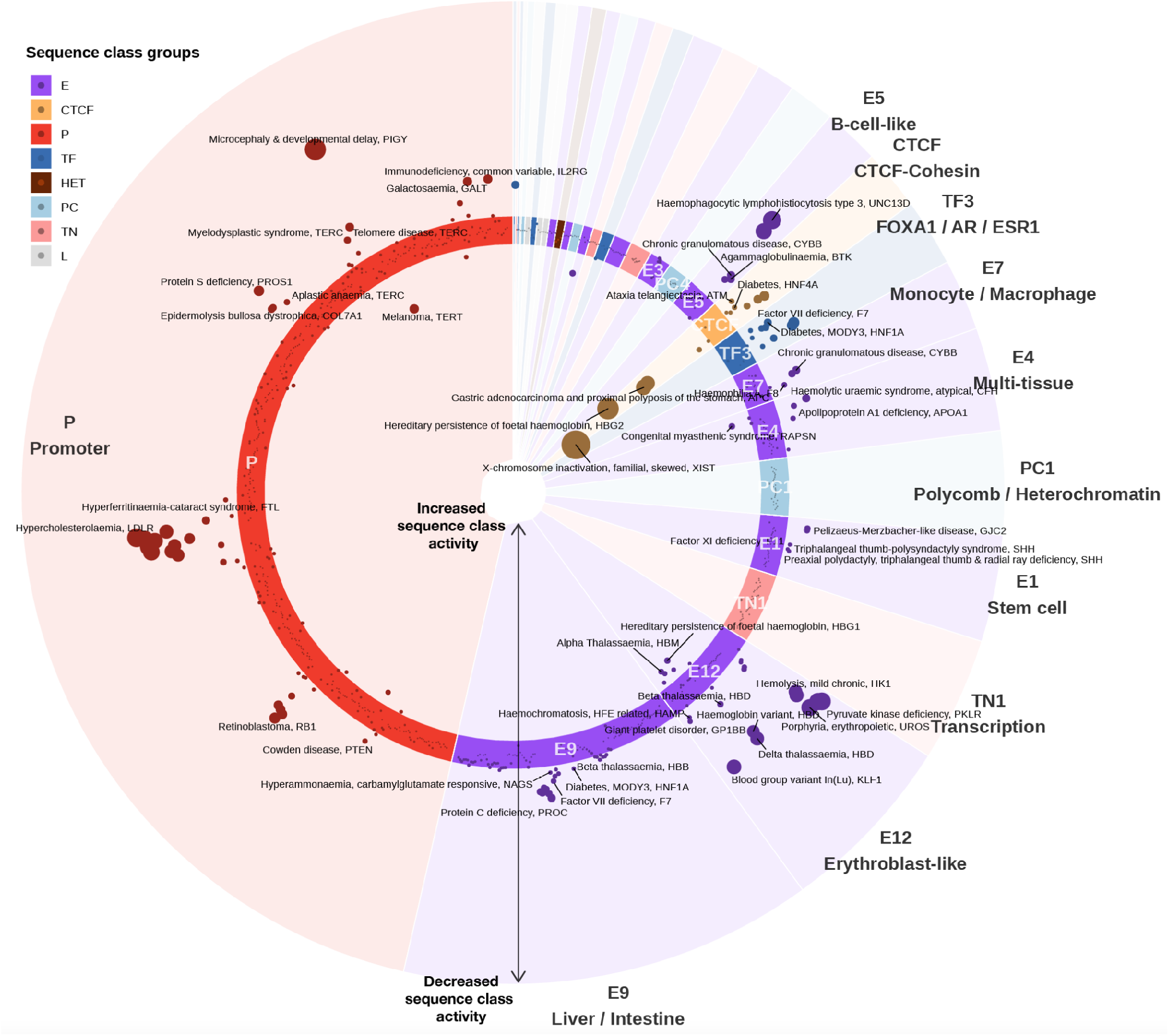
Disease regulatory mutations are predicted to disrupt promoter, CTCF, and tissue-specific enhancer sequence classes. Sequence-class-level mutation effects of pathogenic noncoding HGMD mutations are plotted. A polar coordinate system is used, where the radial coordinate indicates the sequence-class-level effects. Each dot represents a mutation, and mutations inside the circle are predicted to have positive effects (increased activity of sequence class), while mutations outside of the circle are predicted to have negative effects (decreased activity of sequence class). Dot size indicates the absolute value of the effect. Mutations are assigned to sequence classes based on their sequences and predicted effects (Methods). Within each sequence class, mutations are ordered by chromosomal coordinates. The associated disease and gene name are annotated for each mutation, and only the strongest mutation is annotated if there are multiple mutations associated with the same disease, gene, and sequence class.

Importantly, ‘E’ and ‘P’ sequence classes cover almost all classes that explain a high proportion of heritability for GWAS traits and diseases--the same sequence classes inferred to be under strong evolutionary constraints (Figure 3a, Supplementary File 4). We observed three main groups of traits that share similar heritability composition signatures across sequence classes. The first group is blood-related traits, which contains two subgroups of immune-related and non-immune-related traits. The majority of heritability signals in blood-related traits are explained by enhancer classes for the relevant cell type(s), such as monocyte/macrophage enhancer (E7) for *Monocyte Count*, B-cell-like enhancer (E5) for *Auto Immune Traits*, and erythroblast-like (red blood cell progenitor) enhancer (E12) for *Red Blood Cell Distribution Width,* which measures the range of variation in red blood cell volume. Furthermore, autoimmune-related traits are selectively associated with the immune cell type enhancer sequence classes E5 (B-cell like), E11 (T-cell), and E7 (monocyte/macrophage), while erythroblast-like enhancer E12 is specifically linked to red-blood-cell-related traits. Therefore, sequence classes can dissect the cell-type-specific regulatory architecture of traits and diseases with heritability decomposition, even without relying on gene-level information.

Cognitive and mental traits (*Morning Person*, *Neuroticism*, *Smoking Status*, *Years of Education*, *College Education*) have similar sequence-class-level heritability decompositions as well; for this second group of traits, heritability was mostly explained by brain enhancer (E10 and E3) and stem cell enhancer (E1) sequence classes. The link to E1 is consistent with our observation that E1 was also moderately enriched for active enhancer mark H3K4me1 in brain cell types (Figure 2a, Supplementary Figure 7) and is positively correlated with gene expression in brain tissues (Figure 2b).

The third group of traits is intriguingly diverse, including *Balding*, *Lung Forced Vital Capacity*, *Waist-hip Ratio*, *Height*, and *Heel T-score.* The heritability of these traits are mostly explained by multi-tissue enhancer classes (E4, E2, and E8), which show activity in epithelial cells, fibroblast, muscle, and many other cell types. Enhancer activity across multiple tissues in the body may explain the diverse phenotypes that are associated with these traits.

Beyond these three groups, there are a number of traits with unique heritability patterns that are also linked to highly relevant sequence classes. For example, the *High Cholesterol* trait was most associated with the liver and intestine enhancer sequence class (E9), which is consistent with the physiology of cholesterol metabolism and known etiology of this condition^33^. E9 was also linked to red-blood-cell-related traits, in line with the role of liver in erythropoiesis.

Finally, the promoter sequence class P uniquely explained a sizable proportion of heritability in nearly all traits, suggesting a near-universal involvement of promoter sequence variations in all traits and diseases.

We next assessed whether our new sequence classes could explain GWAS heritability beyond that explained by annotations discovered in prior studies. To this end, we performed LDSR analysis with our whole genome annotations of sequence classes conditioned on an up-to-date set of previously identified baseline annotations (v2.2, https://alkesgroup.broadinstitute.org/LDSCORE/). We uncovered 83 significant sequence-class-trait associations with a corrected p-value cutoff of <0.05 (Supplementary File 5). 70% of all UKBB GWAS traits and 9/13 of the E and P sequence classes have at least one significant association after multiple hypothesis testing correction (Supplementary File 5). This finding suggests that sequence classes can identify extensive new regulatory signals that enrich GWAS interpretation.

### Disease mutations are predicted to disrupt the activities of sequence classes

Sequence-class-level effects enable the prediction of specific regulatory mechanisms at the individual, pathogenic mutation level. To showcase our framework’s capability to predict the mechanisms of individual mutations, we used Sei to predict the direction and magnitude of sequence-class-level mutation effects for all 853 regulatory disease mutations from the Human Gene Mutation Database (HGMD)^34^. For systematic classification and quantification of these mutations, we assign each mutation to an affected sequence class based on its mutation effects (the sequence class with the strongest score change) and the sequence that it alters (Methods).

Overall, the average variant effect score of disease mutations is 4.2x larger than the de novo mutations in healthy individuals (0.903 vs 0.217, p<2.2e-16, Wilcoxon rank-sum test two-sided, max absolute effect across sequence classes) and 6.5x larger than the 1000 Genomes common variants with AF>0.01 (0.903 vs 0.139, p<2.2e-16). Here we focus on analyzing the mutations with the strongest predicted effects (>1.1, n=138/853), where predicted effect refers to the variant effect of the assigned sequence class for each mutation (Figure 5, Supplementary Figure 18). Because sequence-class-level variant effects are directional--that is, predicting whether the alternative allele increases or decreases sequence-class level activity--we are able to discover that while the majority (∼80%) of pathogenic mutations with strong predicted effects are predicted to decrease sequence class activity, the remaining 20% of HGMD pathogenic mutations are predicted to increase sequence class activity. Moreover, perturbations to E-, P-, and CTCF-classes make up >99% of the mutations with strong predicted effects on sequence class activity (Supplementary File 6): 44.9% are predicted to affect tissue-specific E sequence classes, 38.4% are predicted to affect the P promoter sequence class, and interestingly, 15.9% are predicted to affect the CTCF-cohesin sequence class (Methods).

We found that almost all mutations with strong predicted effects in cell-type-specific E sequence classes contributed to diseases relevant to that same cell type (Figure 5, Supplementary File 6)-- for most of these mutations, the nearby gene is known to be relevant to the disease but the molecular mechanisms of regulatory disruption is unknown. For example, mutations causing Protein C deficiency and Hemophilia B, two diseases characterized by the deficiency of specific plasma proteins produced in the liver (protein C and coagulation factor IX, respectively), are predicted to decrease E9 liver/intestine sequence class activities. Blood cell-type-specific enhancer sequence classes are disrupted in distinct blood-related diseases and deficiencies relevant to the corresponding cell type: the E12 erythroblast-like enhancer sequence class is disrupted in red blood cell-specific diseases such as pyruvate kinase deficiency, erythropoietic porphyria, delta-thalassemia, and beta-thalassemia; the E7 monocyte/macrophage-like sequence class is disrupted in monocyte and macrophage-related chronic granulomatous disease; and the E5 B-cell-like enhancer sequence class is disrupted in X-linked agammaglobulinemia, a functional deficiency of B-cell. For developmental diseases, such as preaxial polydactyly triphalangeal thumb and radial ray deficiency and triphalangeal thumb-polysyndactyly syndrome, the E1 embryonic stem cell-specific enhancer sequence class is predicted to be disrupted by mutations in a known distal enhancer of Sonic Hedgehog (SHH) (chr7:156583951 G>A^35^, chr7:156583949 G>C^36^), a gene that plays a crucial role in the positioning and growth of limbs, fingers, and toes during development.

In addition, 38% of the regulatory mutations with strong predicted effects affect the activity of the promoter sequence class P, including a hypercholesterolemia mutation near the LDLR gene (chr19:11200089 C>T^37^), a microcephaly & developmental delay mutation near the PIGY gene (chr4:89444948 C>T^38^), and a retinoblastoma mutation near the RB1 gene (chr13:48877851 G>T^39^). The high proportion of mutations perturbing the P sequence class likely reflects both the critical role of promoters in diseases and the emphasis on promoter-proximal mutations in past studies.

While the mutations we’ve discussed thus far are negative effect mutations which decrease sequence class activity, 20% of HGMD pathogenic mutations are predicted to increase sequence class activity. Indeed, these mutations included many known gain-of-function mutations, which validated our predictions. The highest increase in sequence class activity was observed for a mutation (chrX:73072592 G>C) near the XIST gene that skews X-inactivation of the mutant chromosome in females^40^; this mutation was predicted to increase the activity of the CTCF sequence class and has been experimentally validated to increase CTCF binding^41^. Similarly, positive effect predictions for ‘E’ and ‘P’ sequence classes were also validated by previously studied mutations: an alpha-thalassemia mutation near the HBM gene (chr16:209709 T>C^42^) known to create a GATA1 binding site and increase intergenic transcription was predicted to increase the activity of the erythroblast-specific E12 sequence class, and a TERT gene mutation found in individuals with familial melanoma (chr5:1295161 T>G^43^) was predicted to increase the activity of P. Beyond this, many mutations predicted to have strong positive effects were not previously understood. For example, a mutation near the HBG1 gene (chr11:5271262 A>G^44^) that causes persistence of fetal hemoglobin is also predicted to increase the activity of the erythroblast-specific E12 sequence class. Previously, this mutation was known to create an ATGCAAAT octamer^44^ that matches the POU family transcription factor motif, but its functional consequences were unclear.

Notably, even though pathogenic mutations from prior genetics studies are subjected to selection bias, the observation of pathogenic mutations with strong impacts on E-, P-, and CTCF-sequence classes are consistent with our estimation that regulatory sequence classes are under strong evolutionary constraints, and strong disruptions to these classes are likely to cause health consequences. We also note that the pathogenic mutations with strong positive effects on the CTCF class do not contradict the population allele frequency analysis which inferred that further increase of activity on the CTCF sequence class is generally tolerated, because all these pathogenic mutations are located in sequences in other sequence classes, but their sequence class identities are altered to the CTCF class by these mutations. This is in contrast to the allele frequency analysis, which only focuses on variants located in sequences in the CTCF sequence class.

Therefore, sequence-class-level effects both corroborate existing regulatory mechanisms and propose new mechanisms for individual pathogenic mutations. We expect our framework to be a valuable tool in accelerating genetic discoveries of disease-causal mutations and their mechanisms in the regulatory genome.

## Discussion

We developed a genome-wide sequence-based map of regulatory activities using sequence classes, a vocabulary for genomic sequence activities discovered using a data-driven, systematic method. Our deep-learning-based framework uses a compendium covering 21,907 publicly available cis-regulatory profiles and the whole genome sequence to create a mapping from any sequence to a comprehensive set of sequence classes. This provides a global sequence-based view of sequence regulatory activities and allows for the quantitative prediction of variant effects on sequence class activities. Sequence classes are a concise vocabulary of regulatory activities that is interpretable, quantifiable, and easily analyzed globally (across all sequence classes) and individually. To our knowledge, it is the first such attempt to systematically map regulatory activities from any sequence.

We demonstrated that E- and P-sequence classes are strongly enriched in trait and disease GWAS heritability and under evolutionary constraints. Importantly, sequence classes provide insights into the mechanisms of individual pathogenic mutations by predicting effects on the function of tissue-specific enhancers, promoter activity, and long-range genome interactions (e.g. CTCF-cohesin sequence class). Using sequence-class-level variant effect predictions, we linked many pathogenic mutations to tissue-specific regulatory changes in the relevant tissues. These predictions point to potential mechanisms that can be experimentally tested in the future.

Sequence classes leverage a sequence model trained on most publicly available cis-regulatory profile data; however, there remains substantial space for improvement as more data becomes available. For example, we are still lacking data for many cell types, developmental stages, transcription factors, and combinations of chromatin targets measured in new cell types or conditions. More data that covers currently undercharacterized cell types and developmental stages will likely enable the identification of still more cell-type-specific and developmental stage-specific sequence classes, defining sequence classes with increasingly fine-grained regulatory resolution. Furthermore, development of new computational methods to define, for example, hierarchical or combinatorial representations of sequence classes may be needed to make such fine-grained classes easy to interpret and use. Because interpretability and robustness were our major goals in designing sequence classes, we chose to use clustering to generate the sequence classes and a linear projection step to compute corresponding scores. It is conceivable that a more expressive model such as an end-to-end neural network can further improve sequence class predictions, and we expect that increasing the expressiveness of the model while maintaining interpretability and robustness will be an interesting future challenge.

This work demonstrates the potential of sequence classes to discover regulatory disruptions in human diseases, through both the aggregation of genome-wide variant association signals and prediction of the impact of individual mutations. We provide sequence classes and the Sei model as a resource for further research into understanding the regulatory genetic landscape of human health and diseases. Our framework is applicable to any variant, regardless of whether it is common, rare, or never previously observed, and we expect it to be a powerful tool for understanding the mechanistic effects of noncoding mutations in human health.

## Methods

### Training data

21,907 cis-regulatory profiles in peak format were compiled from the processed files of the Cistrome^5^, ENCODE^3^, and Roadmap Epigenomics projects^4^. The Cistrome Project, which systematically processed publicly available cis-regulatory profiles, contributed the majority of the profiles predicted in Sei (19,905). We excluded profiles from Cistrome with less than 1000 peaks. Genome sequences are from the GRCh38/hg38 human reference genome. The full list of cis-regulatory profiles is available in Supplementary File 1.

### Deep learning sequence model training

The Sei model is trained to predict 21,907 transcription factor binding, histone marks, and DNA accessibility from cis-regulatory profile peaks at the center of 4kb sequences.

The model architecture is composed of three sequential sections: 1) a convolutional network with dual linear and nonlinear paths, 2) residual dilated convolution layers, 3) spatial basis function transformation and output layers. A detailed specification of the model is available in Supplementary File 7 and in the code repository (https://github.com/FunctionLab/sei-framework, downloadable from https://doi.org/10.5281/zenodo.4906996). In the convolutional architecture, we introduced a new design composed of both linear and nonlinear convolution blocks. The nonlinear blocks are composed of convolution layers and rectified linear activation functions (ReLU), similar to regular convolutional networks. The linear blocks have the same structure as the nonlinear blocks but do not include activation functions to facilitate learning of linear dependencies. Each nonlinear block is stacked on top of a linear block with a residual connection adding the input of the nonlinear block to the output, allowing the computation to go through either the linear or nonlinear path. Dilated convolutional layers with residual connections further expands the receptive fields without reducing spatial resolution. Finally, spatial basis functions are used to reduce dimensionality of the spatial dimension while preserving the capability to discriminate spatial patterns of sequence representations. Specifically, in the Sei model, a B-spline basis matrix (256×16) with 16 degrees-of-freedom across 256 uniformly-spaced spatial bins is generated and multiplied with the convolutional layers output to reduce the 256 spatial dimensions to 16 spline basis function dimensions. After the spline basis function transformation, a fully-connected layer and an output layer are used for integrating information across the whole sequence and generating the final 21,907-dimensional predictions.

Training, validation, and testing datasets are specified by different sets of chromosomes in the hg38 genome (holding out chromosome 8 and 9 for the test set and chromosome 10 for the validation set), and samples drawn uniformly across the hg38 genome for these partitions, excluding regions specified in the ENCODE blacklist^45^. For training, we sampled training sequences and their labels on-the-fly from the training set of chromosomes using Selene^46^. As a result, almost all training samples are drawn from unique genomic intervals with distinct start and end positions to reduce overfitting during the training process. For each 4kb region, a 21,907-dimensional binary label vector is created for the 21,907 cis-regulatory profiles based on whether the center basepair overlaps with a peak in each of the profiles. The model is implemented in PyTorch and trained with Selene. A detailed training configuration file is available at https://github.com/FunctionLab/sei-framework/blob/main/train/train.yml.

### Model performance

We computed the AUROC and AUPRC for all cis-regulatory profiles predicted by Sei on the test holdout dataset, excluding profiles that had fewer than 25 positive samples in the test set. Additionally, to assess the correlation structure of the predictions, we compared the rank-transformed pairwise Spearman’s rank correlations for the predicted cis-regulatory profiles to the pairwise correlations for the true labels (peak calls provided in Cistrome DB).

The model performance comparison between DeepSEA and Sei is computed on the 2,002 cis-regulatory profiles from Roadmap and ENCODE that both DeepSEA and Sei predict. Because both models have the same chromosomal test holdout (chr8 and chr9), we use the regions specified in the DeepSEA test holdout set to create a common test dataset of sequences and labels on which to evaluate the models.

### Sequence classes

We selected 30 million genomic positions that uniformly tile the genome with 100bp step size and then computed Sei predictions for 4kb sequences centered at each position. Sequences overlapping with ENCODE blacklist regions^45^ or assembly gaps (“N”s) are removed. To process the 30 million x 21,907 predictions matrix, the dimensionality is first reduced with principal component analysis (PCA). The PCA transformations were fitted with incremental PCA using a batch size of 1,000,000 for one pass of the whole dataset, and genomic positions were randomly assigned to batches. The top 180 principal components, scaled to unit variance, were used for constructing a nearest neighbor graph where each node is connected to its k-nearest neighbors by Euclidean distance (k=14). Louvain community clustering with default parameters was applied to the nearest neighbor graph with the python-louvain package, which resulted in 61 clusters. We refer to the largest 40 clusters as sequence classes and exclude the remaining (smallest) 21 clusters, which constitute <2.6% of the genome, from our analyses due to their size. These 21 clusters mainly display Low signal or Heterochromatin like enrichment (Supplementary Figure 19). We refer to this cluster assignment to sequence classes at 100bp resolution as sequence class annotations. We visualized the genome-wide predictions by computing UMAP embedding with a subsample of PCA-transformed Sei predictions of 30 million sequences, and then fine-tuned the visualization with OpenTSNE. The detailed procedures are available in our code repository (https://github.com/FunctionLab/sei-manuscript).

### Sequence class scores

Each sequence class is represented as a unit vector in the 21,907-dimensional cis-regulatory profile space, in the direction of the average prediction of all sequences assigned to this sequence class among the 30 million. In more formal notation, the vector for sequence class *i* is *ν_i_* = 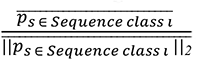, where *p*_s_ represents the 21,907-dimensional Sei prediction for sequence *s*. Each Sei prediction can then be projected onto any sequence class vector to obtain a sequence class-level representation of the prediction, which we call sequence class score or *score_s,i_* = *p*_s_·*ν_i_^T^*. In addition, predicted sequence-class-level variant effects are represented by the difference between the sequence class scores of the sequences carrying the reference allele and the alternative allele, or *score_ν,i_* = *score_alt,i_* – *score_ref,i_*. To better represent predicted variant effects on histone marks, it is necessary to normalize for the nucleosome occupancy (e.g. loss-of-function mutation near TSS can decrease H3K4me3 modification level while increasing nucleosome occupancy, resulting in an overall increase in observed H3K4me3 quantity). Therefore, for variant effect computation, we use the sum of all histone profile predictions as an approximation to nucleosome occupancy and adjust all histone mark predictions to remove the impact of nucleosome occupancy change (non-histone mark predictions are unchanged):

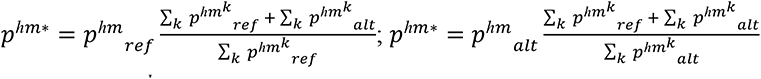

where 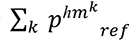 represents the sum over all histone mark predictions (among 21907-dimensions of a prediction) for the reference allele. We generally exclude Low Signal sequence classes in sequence-class-level variant effect analyses because they lack an intuitive biological interpretation.

### Sequence class enrichment of chromatin profiles and genome annotations

We computed the log fold change enrichment of various chromatin profiles and genome annotations for each sequence class based on sequence class annotations (described above, see ‘Sequence classes’). Log fold change enrichment is computed by taking the log ratio of the proportion of a sequence class intersecting with the annotation versus the background proportion of the annotation, where we consider all regions assigned to any sequence class. We computed enrichment for all 21,907 profiles predicted by Sei, filtered the chromatin profiles for each sequence class to only those having Benjamini-Hochberg corrected p-values (Fisher’s exact test, two-sided) below 2.2e-16, and selected the top 25 profiles based on log fold change enrichment. Cistrome Project profile enrichment is computed over 2 million random genomic positions.

The annotation of centromere repeats is obtained from the UCSC RepeatMasker track, and annotations of histone marks over multiple cell types are obtained from the Roadmap Epigenomics project--enrichments for both of these sets of annotations are computed over the entire genome. In addition, we obtained ChromHMM chromatin states from ENCODE ^3^ and tissue and cell-type-specific DHS vocabulary from ^25^.

### Enhancer sequence class correlations with cell-type-specific gene expression

Tissue expression profiles are from GTEx^26^, Roadmap Epigenomics^4^, and ENCODE^3^ and transformed to log-scale RPKM (reads per kilobase per million reads mapped) scores as previously described^13^ and normalized by tissue-average. Specifically, a pseudocount was added before log transformation (0.0001 for GTEx tissues, which are averaged across individuals, and 0.01 for Roadmap and ENCODE tissues). After log transformation, the average scores across tissues were subtracted for each gene; as a result, the processed scores represent log fold change relative to tissue-average.

Gene-wide expression prediction is evaluated on sequence class annotations (from Louvain community clustering) for positions within +/-10kb of the TSSs for these genes. For each enhancer sequence class and tissue, we compute the Spearman correlation between the sequence class annotation coverage and gene expression.

### Correlation between regulatory sequence class variant effects and directional eQTL variant effect sizes

We collected the eQTLs within +/-5kb of gene TSSs from GTEx v8, combined across all GTEx tissues, and computed the Spearman correlation between the top 15k variant effect predictions for each sequence class and the eQTL variant effect sizes (averaged across multiple tissues if the variant is an eQTL in multiple tissues). The p-values are derived from the Spearman’s rank correlation test (two-sided) and BH correction is applied. Low Signal and Heterochromatin sequence classes are excluded from this analysis due to lack of interpretation for their variant effect scores in this context.

Additionally, we collected fine-mapped GTEx eQTLs from eQTL Catalogue^27^ and obtained sequence class scores for eQTLs with posterior inclusion probability > 0.95. Variants are assigned to sequence classes based on the sequence class annotation for the reference genome (i.e. variants are not further selected based on variant effect predictions). For each sequence class, we computed the Spearman correlation between the sequence class scores and the eQTL variant effect sizes in the same way we describe above.

### Evolutionary constraints on variant effects

We computed sequence-class-level variant effects for all 1000 Genomes project phase 3 variants^28^. Variants are assigned to sequence classes based on the 100bp resolution genome-wide assignment derived from Louvain community clustering as described above. For each sequence class we divide variants into 6 bins based on their effects in the same sequence class as illustrated in Figure 3, and summarize common variant (AF>0.01) frequencies in each bin by mean and standard error of the mean. We also estimated statistical significance of allele frequency dependency on sequence-class-level variant effects. For each sequence class, we applied logistic regression separately for positive effect and negative effect variants, to predict common variants (AF>0.01) from the absolute value of sequence-class-level variant effect score, and obtained the significance z-score of the regression coefficient of variant effect. The bidirectional evolutionary constraint z-score is defined as the negative value of the combined z-scores from positive and negative effect variants with Stouffer’s method.

### Partitioning GWAS heritability by sequence classes

UKBB GWAS summary statistics were obtained from ^18^. To study the association of sequence class genome annotation and sequence class variant effects and trait heritability, we performed partitioned heritability LD score regression (LDSR) as described in ^29^. To partition the heritability as sums of heritability explained by each sequence class, we run LDSR with only sequence class annotations and a baseline all-ones annotation. We obtained the estimated proportion of *h*^2^ explained by each sequence class and its standard error with LDSR as implemented in https://github.com/bulik/ldsc. As the estimated proportions can have high variance or even be negative (the true value of heritability explained can only be non-negative), we use a robust and conservative estimator which is the estimated proportion of *h*^2^ subtracted by one standard error, then lower-bounded by zero (the standard error of the estimated proportion of *h*^2^explained is given by LDSR and estimated with the block jackknife procedure as described in^29^).

To assess the contribution of sequence classes to explaining additional heritability when conditioned on known baseline annotations, we also run LDSR with the baseline annotations (v2.2, https://alkesgroup.broadinstitute.org/LDSCORE/). The p-values are derived from the coefficient z-score, and BH correction is applied.

### Sequence class-level variant effect analysis of noncoding pathogenic mutations

We obtained all mutations assigned “DM” and “regulatory” annotation in the Human Gene Mutation Database (HGMD) database (2019.1 release). RMRP gene mutations are excluded because they are likely pathogenic due to impacting RNA function instead of regulatory perturbations, despite being annotated to the regulatory category in HGMD. For every mutation, we predicted the sequence class scores for both the reference and the alternative allele and computed the sequence-class-level variant effect as the predicted scores for the alternative allele subtracting the scores for the reference allele. To provide an overview of sequence-class level effects of human noncoding pathogenic mutations, mutations are first assigned to sequence classes based on the sequence class annotations of the mutation position. For mutations with a strong effect in a different sequence class than the originally assigned sequence class (absolute value higher than the original sequence class by >1 absolute difference and >2.5 fold relative difference), we reassign the mutation to the sequence class with the strongest effects.

## Code and data availability

The Sei framework code is provided in https://github.com/FunctionLab/sei-framework, and the model and associated data files downloadable by following the instructions in the GitHub repository. Code and data for the manuscript results are available at https://github.com/FunctionLab/sei-manuscript.

## Acknowledgements

The authors acknowledge all members of the Troyanskaya lab for helpful discussions. This work was performed using the high-performance computing resources, supported by the Scientific Computing Core, at the Flatiron Institute and the Terascale Infrastructure for Groundbreaking Research in Science and Engineering high-performance computer center at Princeton University. K.M.C. is supported by the National Science Foundation Graduate Research Fellowship Program (NSF-GRFP). O.G.T. is supported by National Institutes of Health grant nos. R01HG005998, U54HL117798 and R01GM071966, U.S. Department of Health and Human Services grant no. HHSN272201000054C and Simons Foundation grant no. 395506. O.G.T. is a senior fellow of the Genetic Networks program of the Canadian Institute for Advanced Research. J.Z. is supported by the Cancer Prevention and Research Institute of Texas grant (RR190071), National Institutes of Health grant DP2GM146336, and the UT Southwestern Endowed Scholars program.

## Author Contributions

K.M.C. and J.Z. conceived the Sei framework, developed the computational methods, and performed the analyses. A.K.W. developed the Sei web server. K.M.C., J.Z., and O.G.T. wrote the manuscript.

---

**Supplementary Figure 1.**
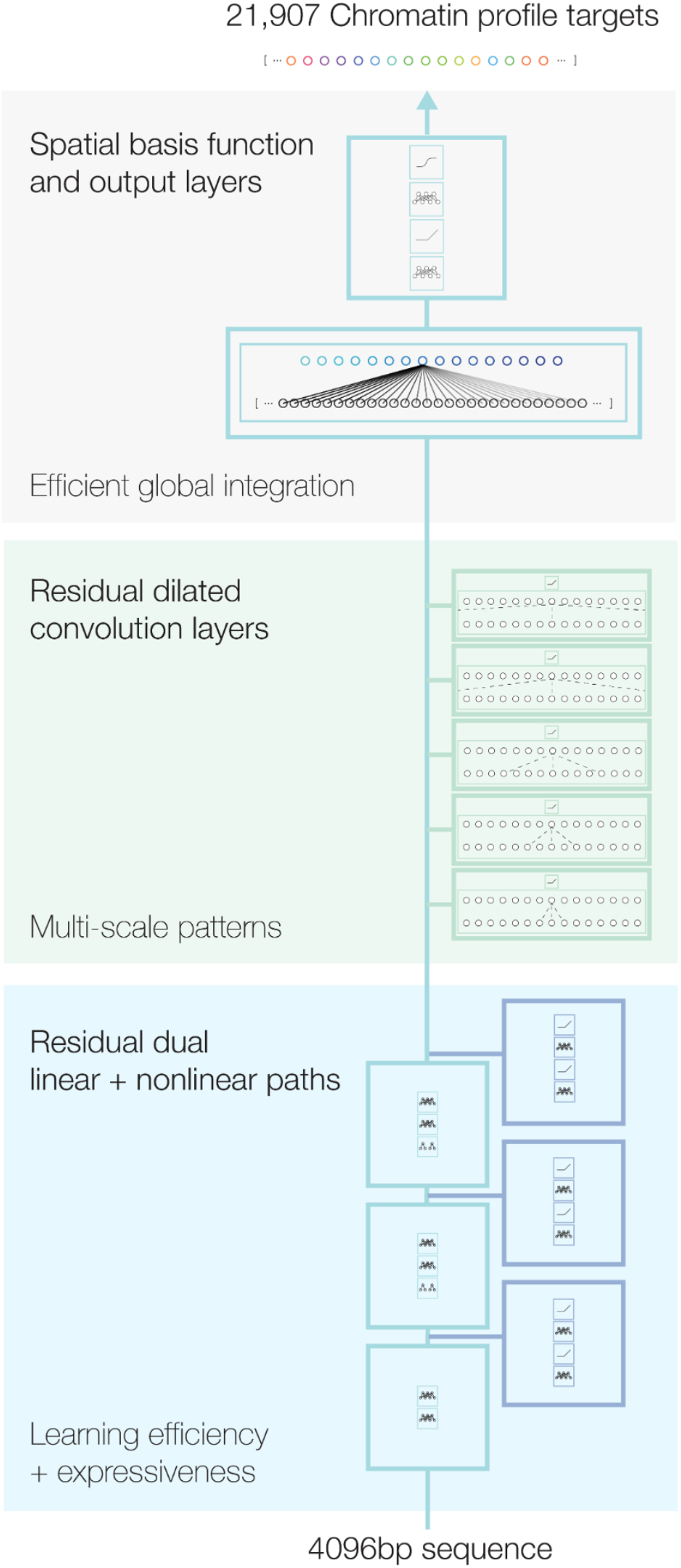
Schematic overview of Sei model architecture. 4096bp sequences, one-hot encoded, are the input to the model (bottom) and the predicted 21,907 cis-regulatory profiles are the output (top).

**Supplementary Figure 2.**
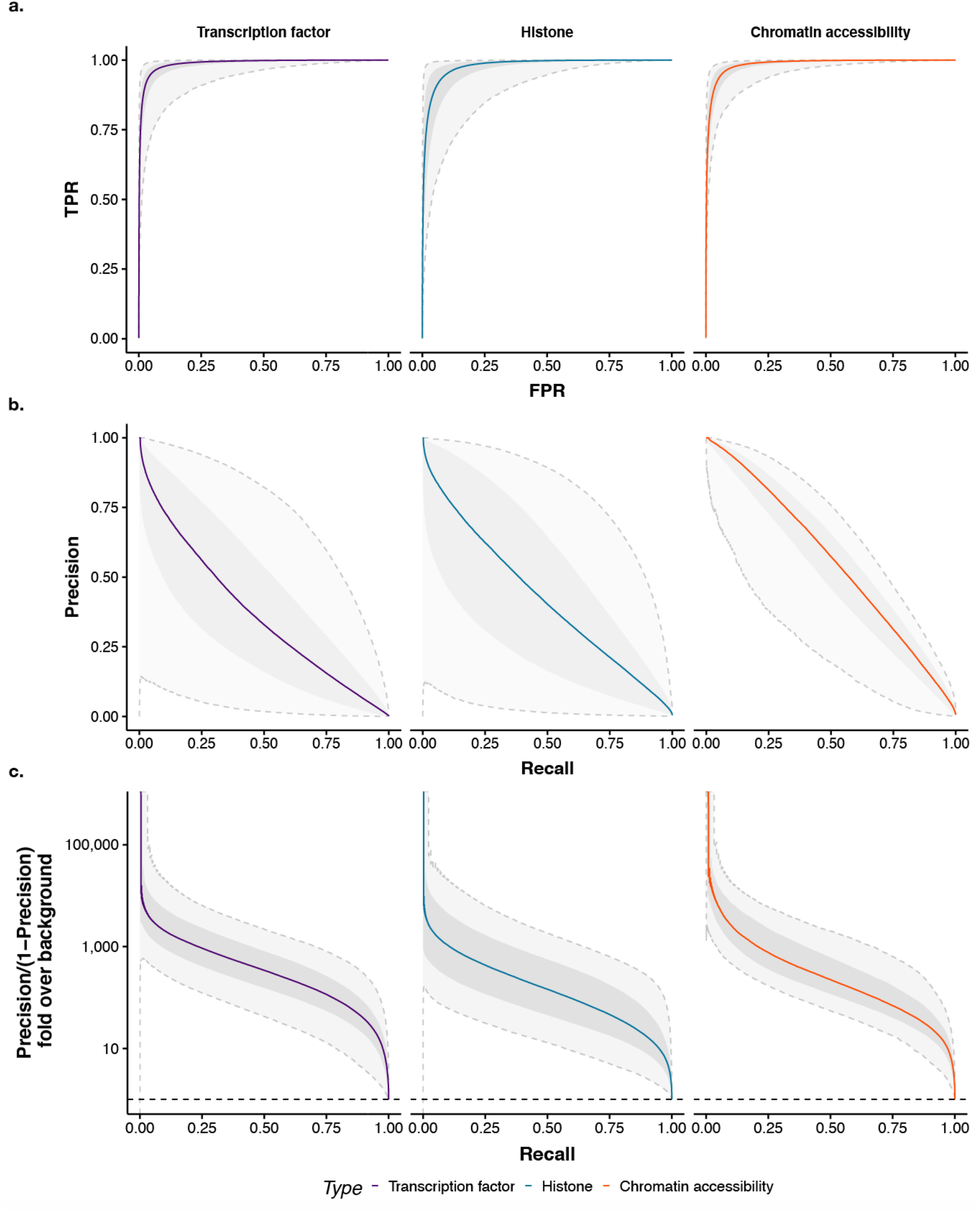
Sei model performance on predicting 21907 cis-regulatory profiles on holdout chromosomes.

**Supplementary Figure 3.**
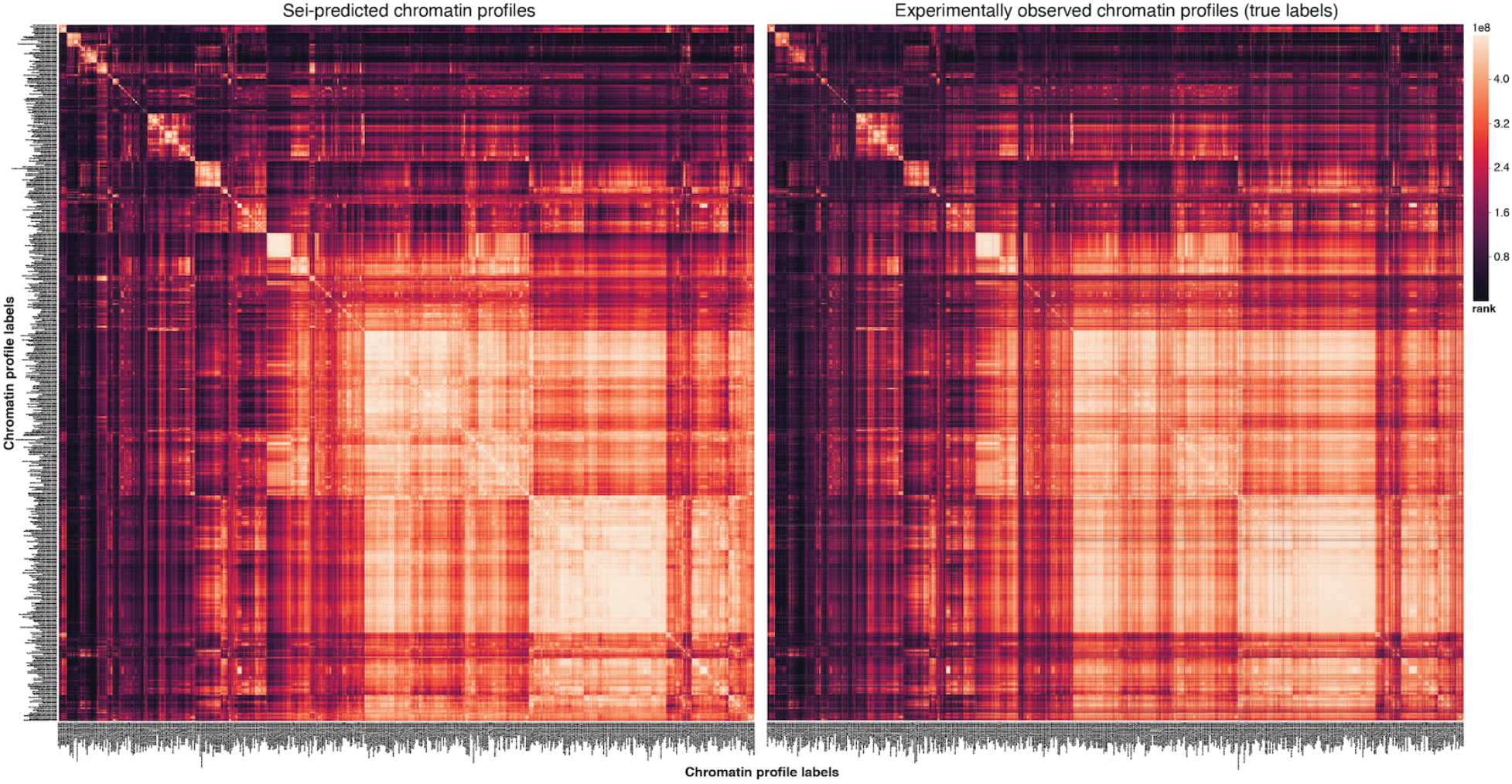
Visualizing the rank-transform of pairwise Spearman correlations for the 21,907 cis-regulatory profiles in Sei. Sei model predictions share a highly similar correlation structure with the experimental observations.

**Supplementary Figure 4.**
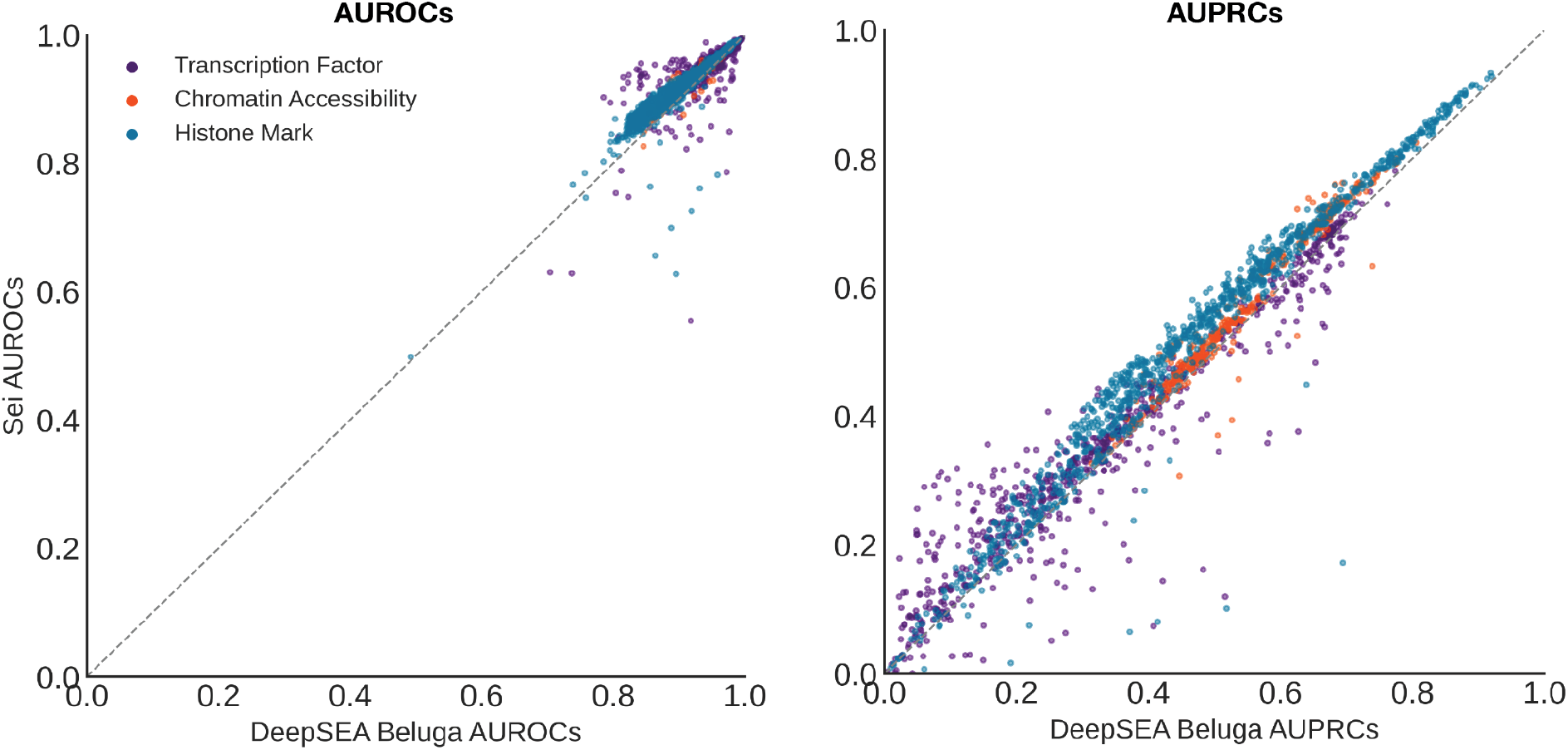
Sei model performance comparison with DeepSEA. Performance on the shared 2002 DeepSEA “Beluga” (2018) cis-regulatory profiles are compared.

**Supplementary Figure 5.**
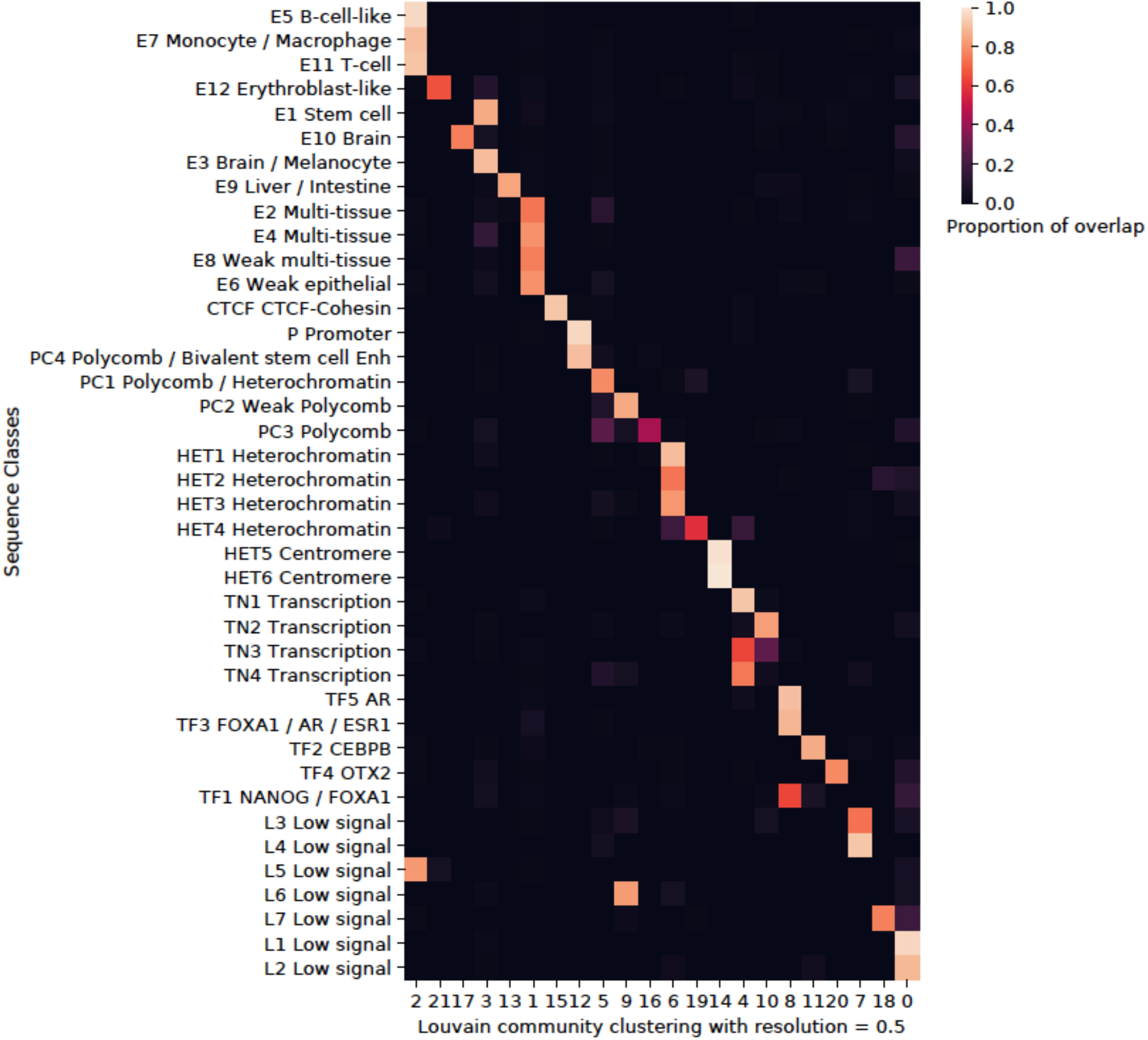
Comparison of sequence classes and Louvain community clustering with resolution = 0.5. For each sequence class, the proportion overlap was computed between sequence classes and a lower resolution clustering for Louvain community clustering. The lower resolution clustering is largely consistent with the original sequence classes, with some clusters combining several related enhancer sequence classes into one.

**Supplementary Figure 6.**
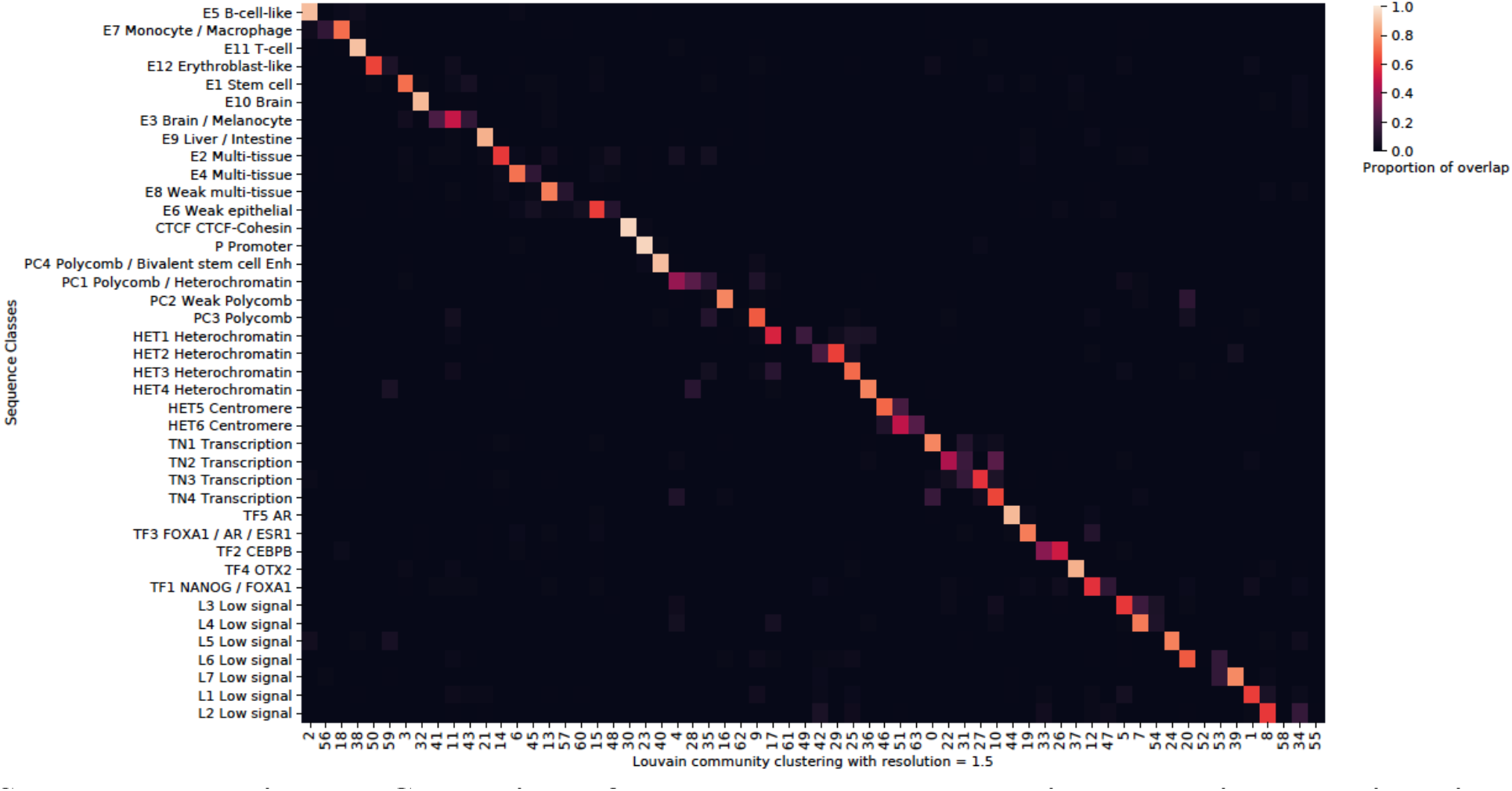
Comparison of sequence classes and Louvain community clustering with resolution = 1.5. For each sequence class, the proportion overlap was computed between sequence classes and a higher resolution clustering for Louvain community clustering. The higher resolution clustering closely resembles the current sequence class clusters.

**Supplementary Figure 7.**
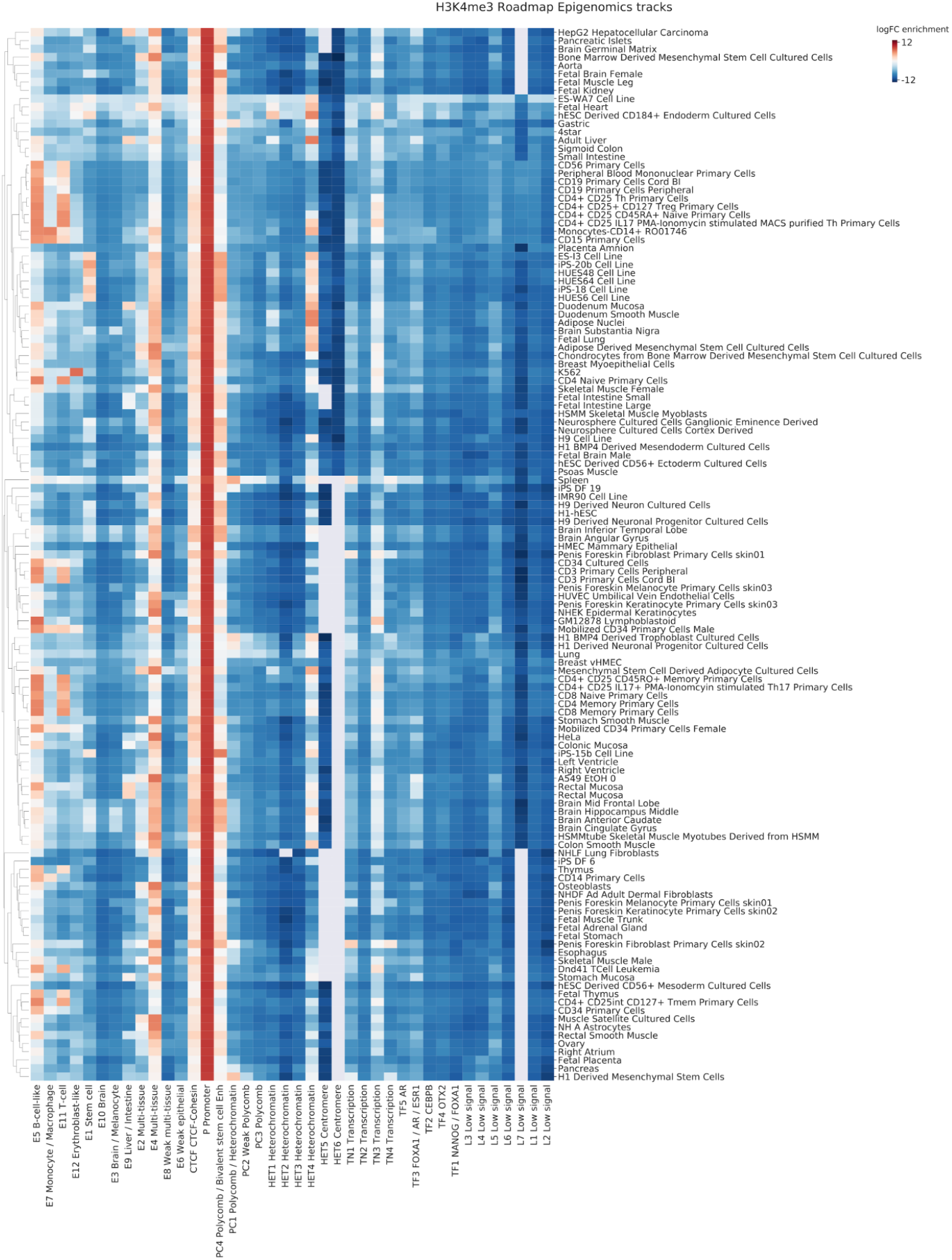
Enrichment of tissue/cell type-specific H3K4me3 (promoter mark) profiles in sequence classes. Log fold change enrichment over genome-average background is shown in the heatmap. No overlap is indicated by the gray color in the heatmap.

**Supplementary Figure 8.**
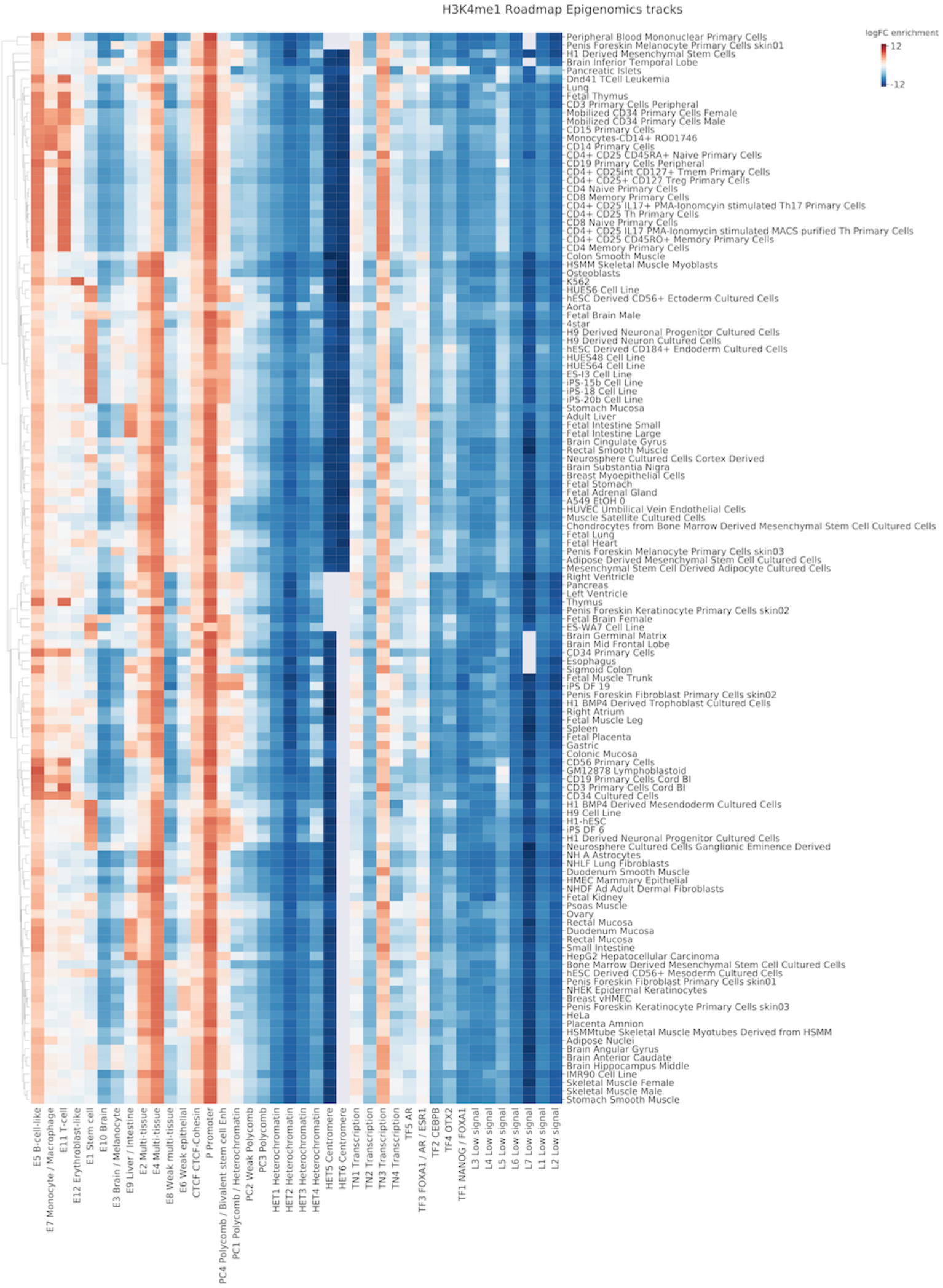
Enrichment of tissue/cell type-specific H3K4me1 (enhancer mark) profiles in sequence classes. Log fold change enrichment over genome-average background is shown in the heatmap. No overlap is indicated by the gray color in the heatmap.

**Supplementary Figure 9.**
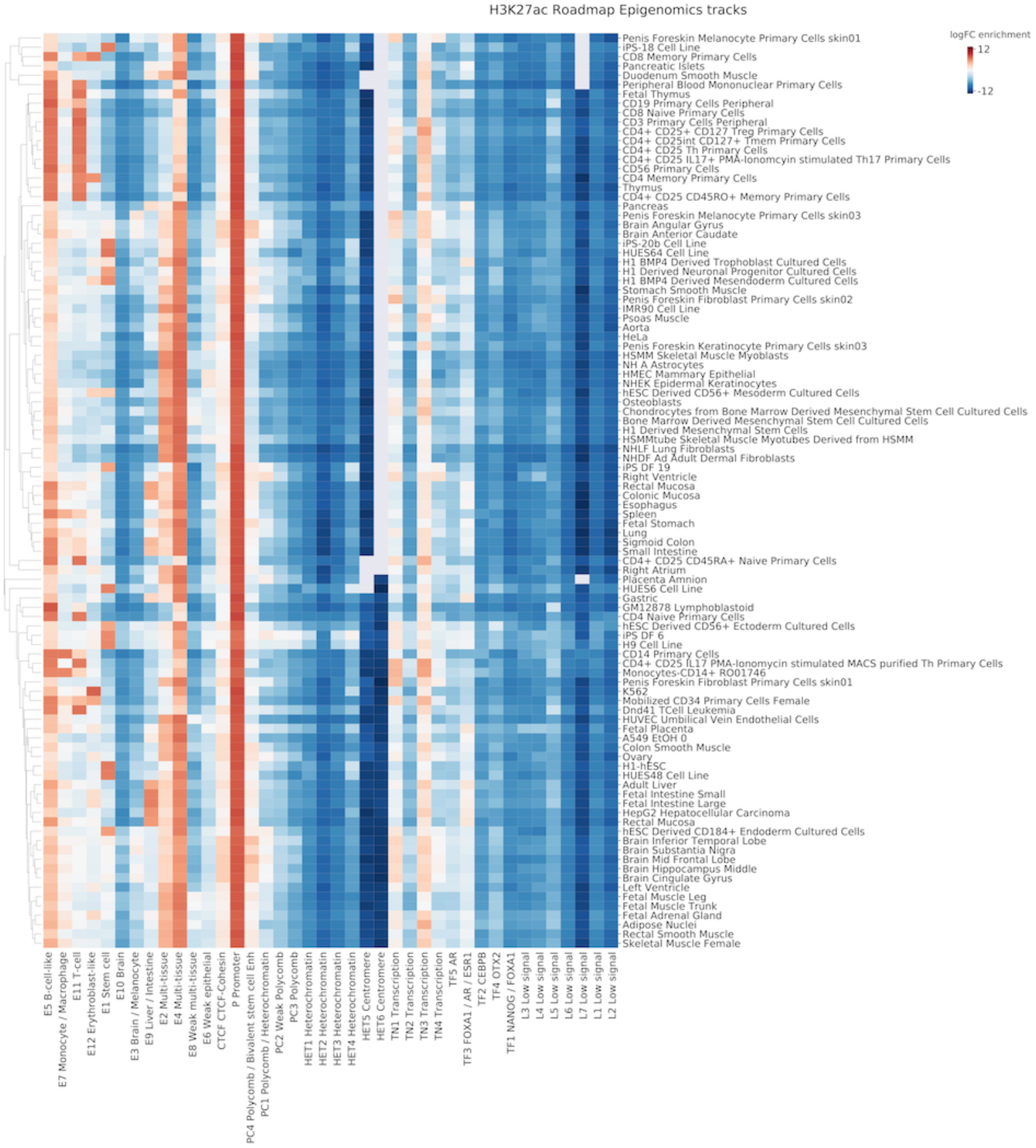
Enrichment of tissue/cell type-specific H3K27ac (enhancer mark) profiles in sequence classes. Log fold change enrichment over genome-average background is shown in the heatmap. No overlap is indicated by the gray color in the heatmap.

**Supplementary Figure 10.**
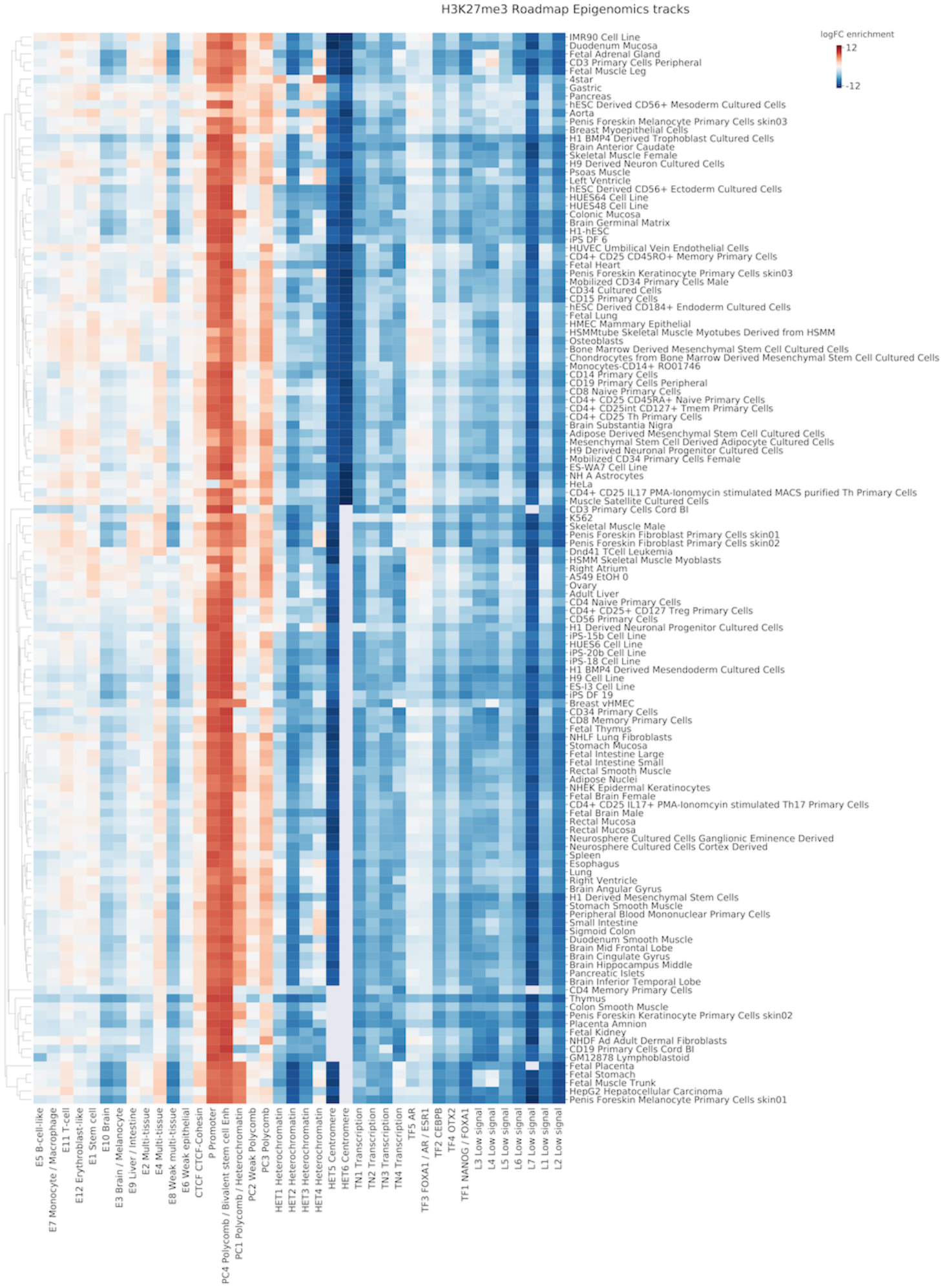
Enrichment of tissue/cell type-specific H3K27me3 (Polycomb mark) profiles in sequence classes. Log fold change enrichment over genome-average background is shown in the heatmap. No overlap is indicated by the gray color in the heatmap.

**Supplementary Figure 11.**
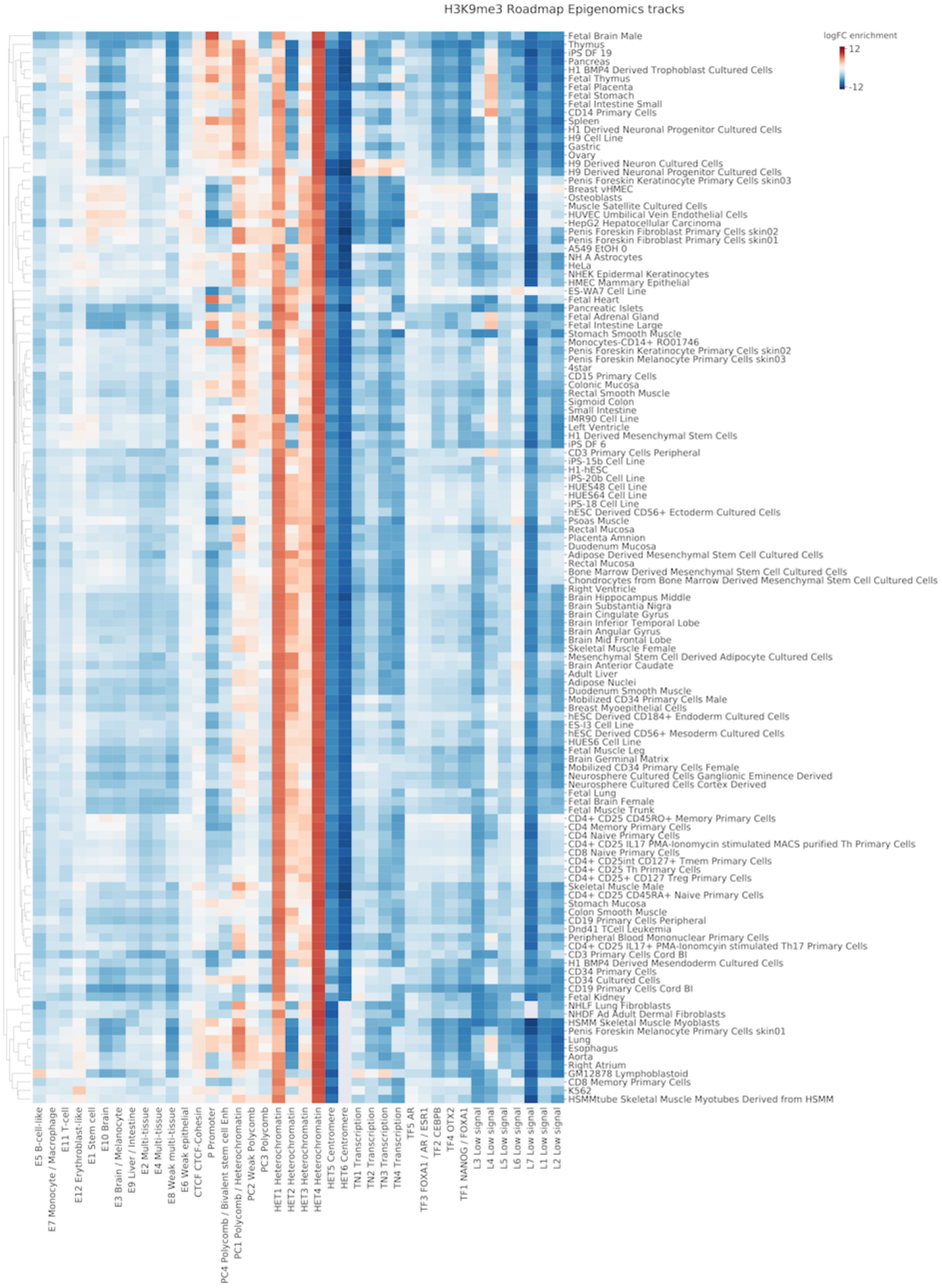
Enrichment of tissue/cell type-specific H3K9me3 (heterochromatin mark) profiles in sequence classes. Log fold change enrichment over genome-average background is shown in the heatmap. No overlap is indicated by the gray color in the heatmap.

**Supplementary Figure 12.**
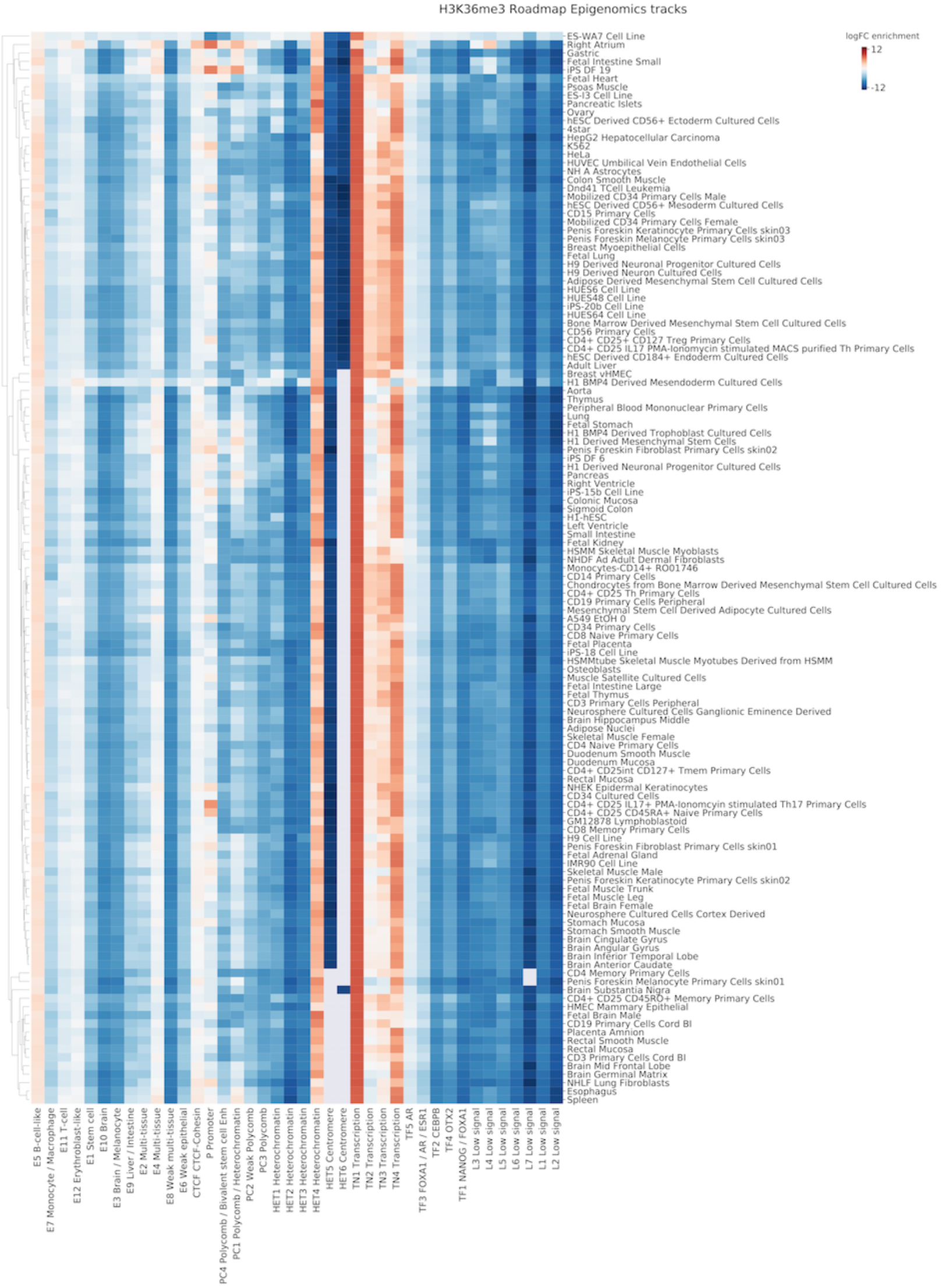
Enrichment of tissue/cell type-specific H3K36me3 (transcription mark) profiles in sequence classes. Log fold change enrichment over genome-average background is shown in the heatmap. No overlap is indicated by the gray color in the heatmap.

**Supplementary Figure 13.**
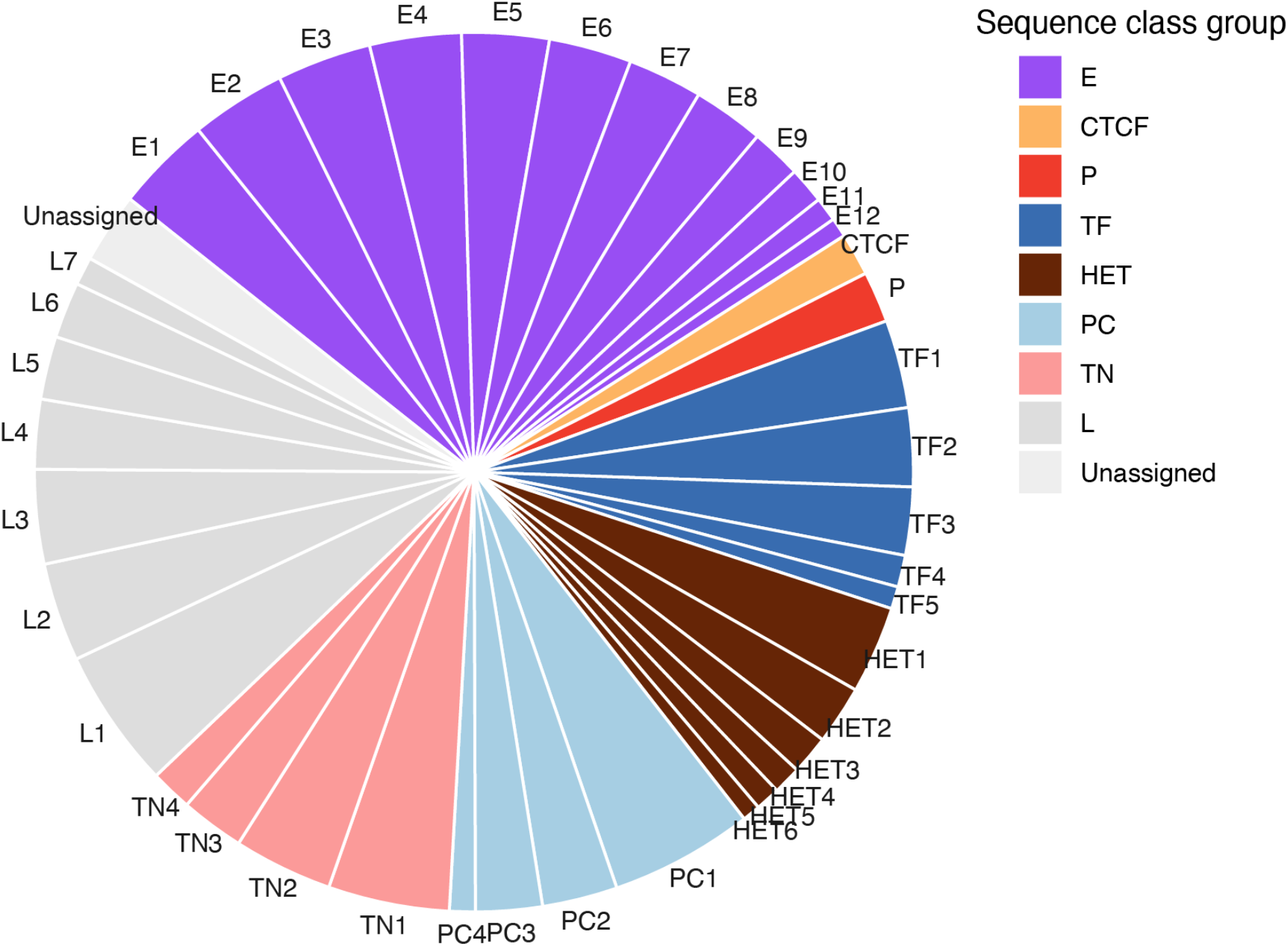
Genome sequence proportion covered by each sequence class. The proportion of each sequence class is shown in the pie chart. Genome-wide sequence class assignments were based on Louvain clustering of Sei predictions of sequence tiling the genome with 100bp step size. The clusters unassigned to sequence classes due to the small size (below top 40 clusters) were categorized as “Unassigned”.

**Supplementary Figure 14.**
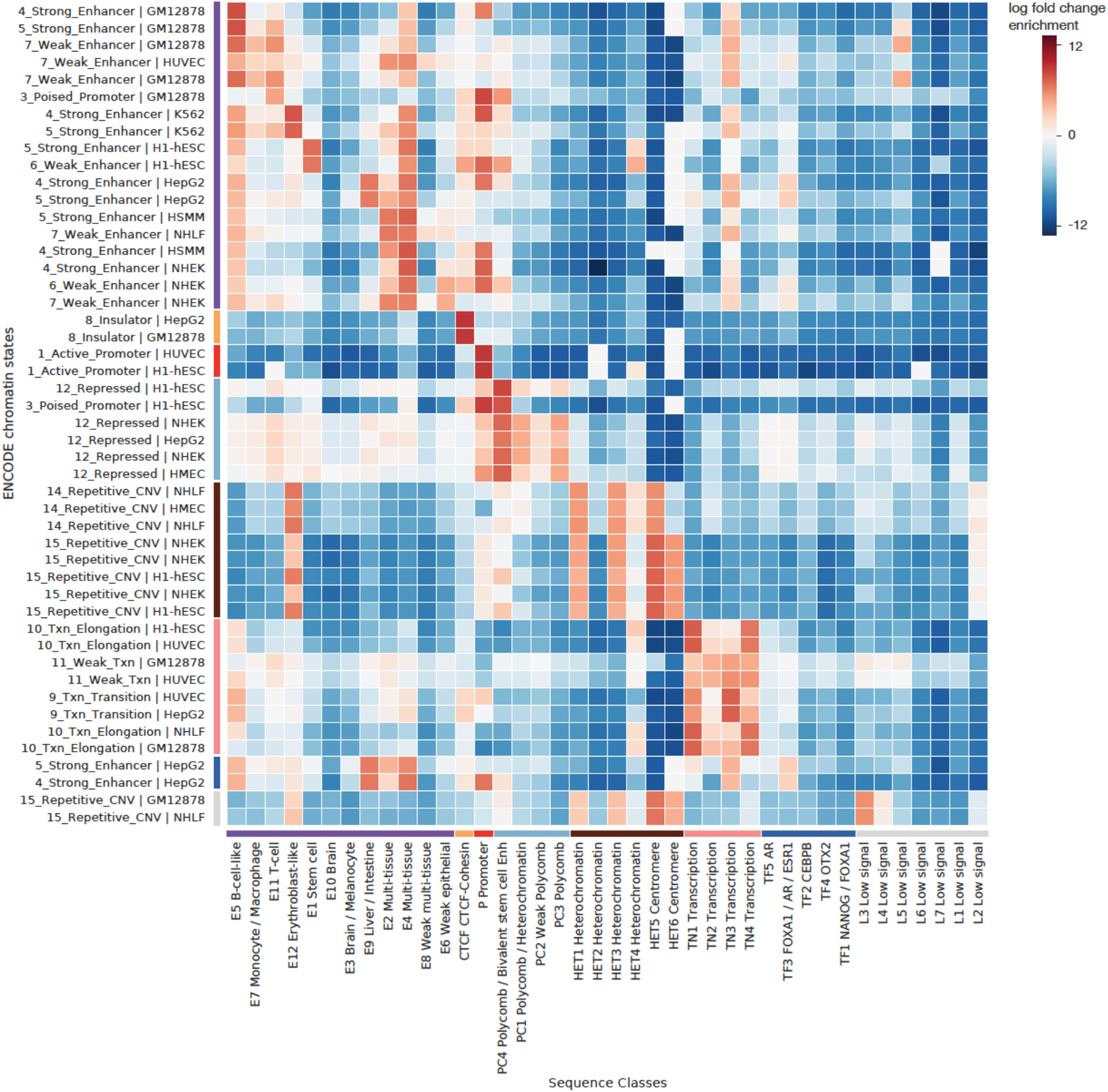
Sequence-class-specific enrichment of ENCODE chromatin states. Log fold change enrichment over genome-average background is shown in the heatmap. Top 2 chromatin states enriched were selected for each sequence class.

**Supplementary Figure 15.**
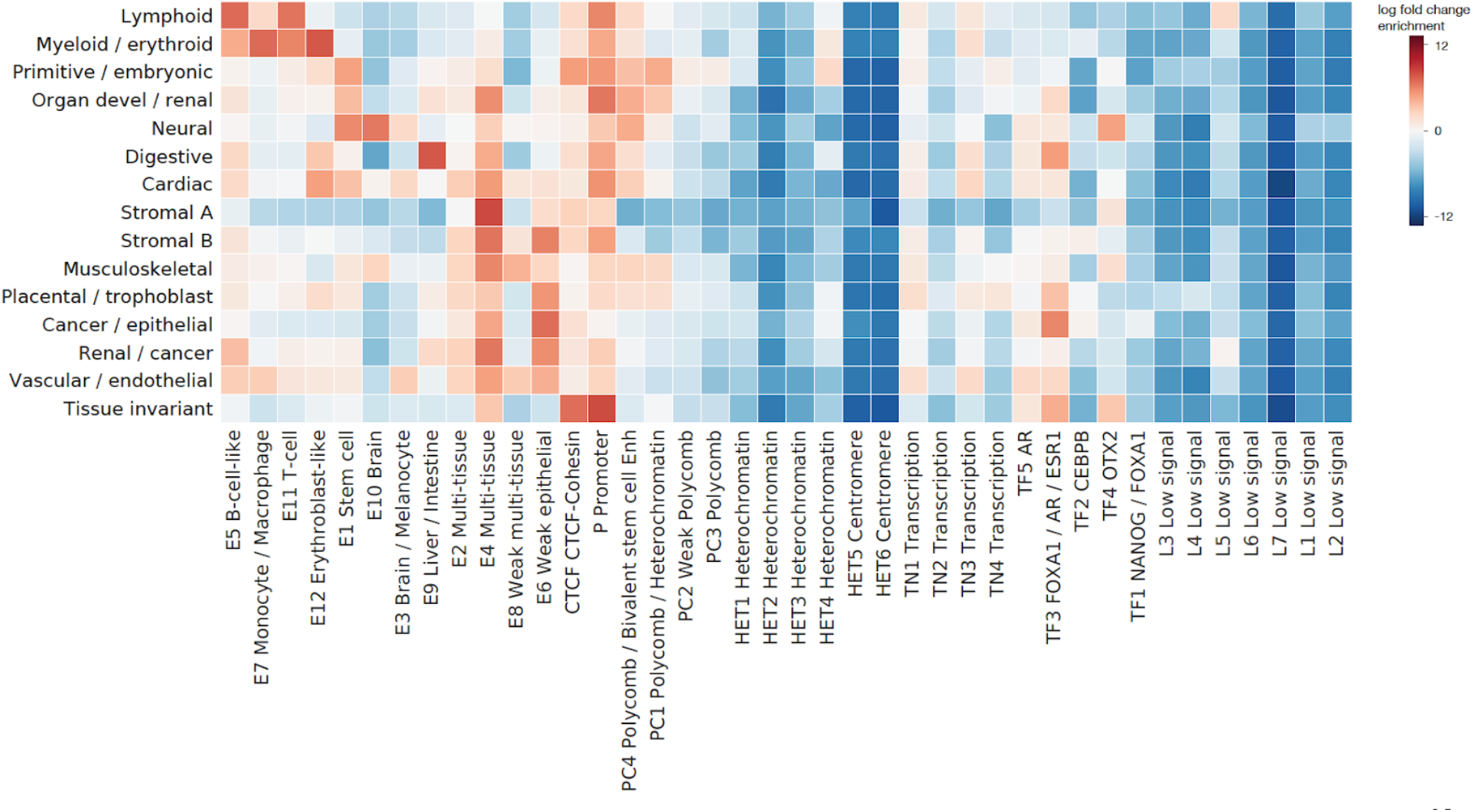
Sequence-class-specific enrichment of tissue-specific DHS vocabulary ^25^. Log fold change enrichment over genome-average background is shown in the heatmap.

**Supplementary Figure 16.**
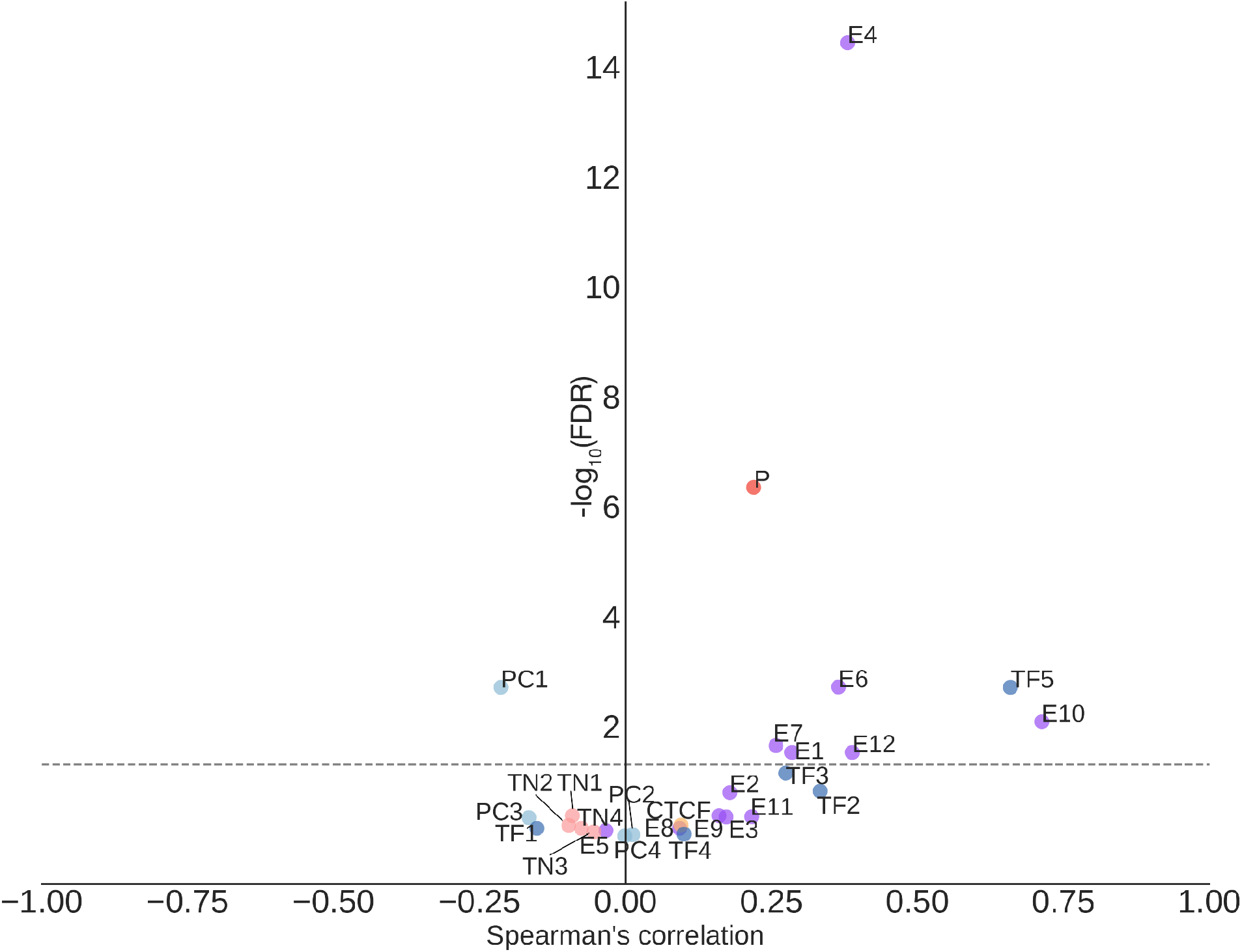
Regulatory sequence-class-level variant effects for SNPs with PIP > 0.95 are predictive of directional GTEx variant gene expression effects. Variants assigned to sequence classes based on the sequence class annotation for the reference genome. The x-axis shows Spearman correlations between the predicted sequence-class-level variant effects and the signed GTEx variant effect sizes (slopes) and the y-axis shows the corresponding log10 p-values. The dotted gray line denotes the Benjamini-Hochberg FDR < 0.05 threshold.

**Supplementary Figure 17.**
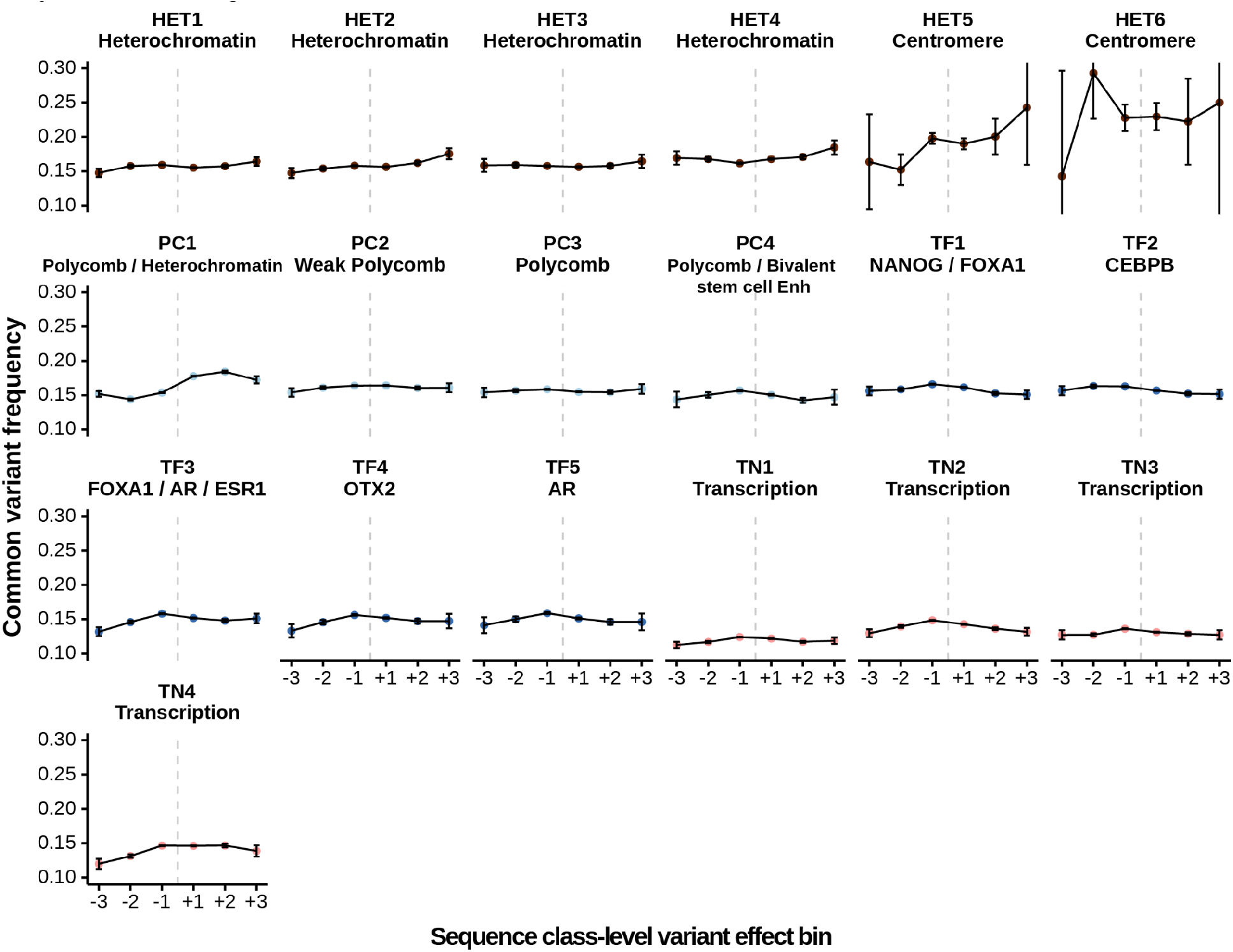
Population allele frequency profiles for variants in heterochromatin, low signal, polycomb, and transcription sequence classes. Comparison of common variant frequencies of 1000 Genomes variants assigned to different sequence classes and variant effect bins. The common variant threshold is >0.01 allele frequency across the 1000 Genomes population. Error bars show +/- 1 standard error(SE). The sequence-class-level variant effects are assigned to 6 bins (+3: top 1% positive, +2: top 1%-10% positive, +1, top 10% -100% positive, -3: top 1% negative, -2: top 1%-10% negative, -1, top 10% -100% negative).

**Supplementary Figure 18.**
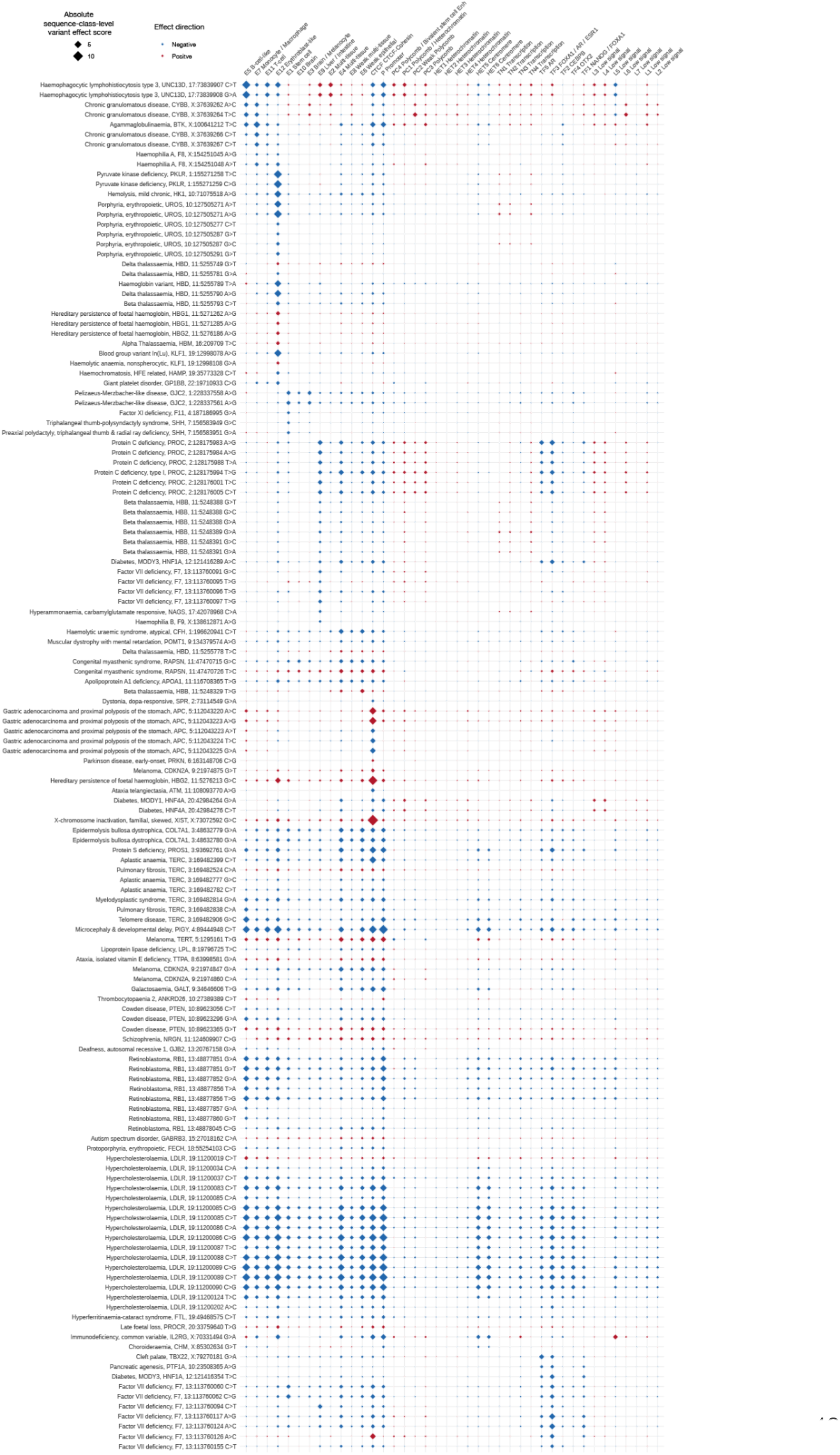
Predicted sequence class-level variant effects for HGMD regulatory disease mutations. HGMD regulatory disease mutations with sequence-class level variant effect score >1.1 are included.

**Supplementary Figure 19.**
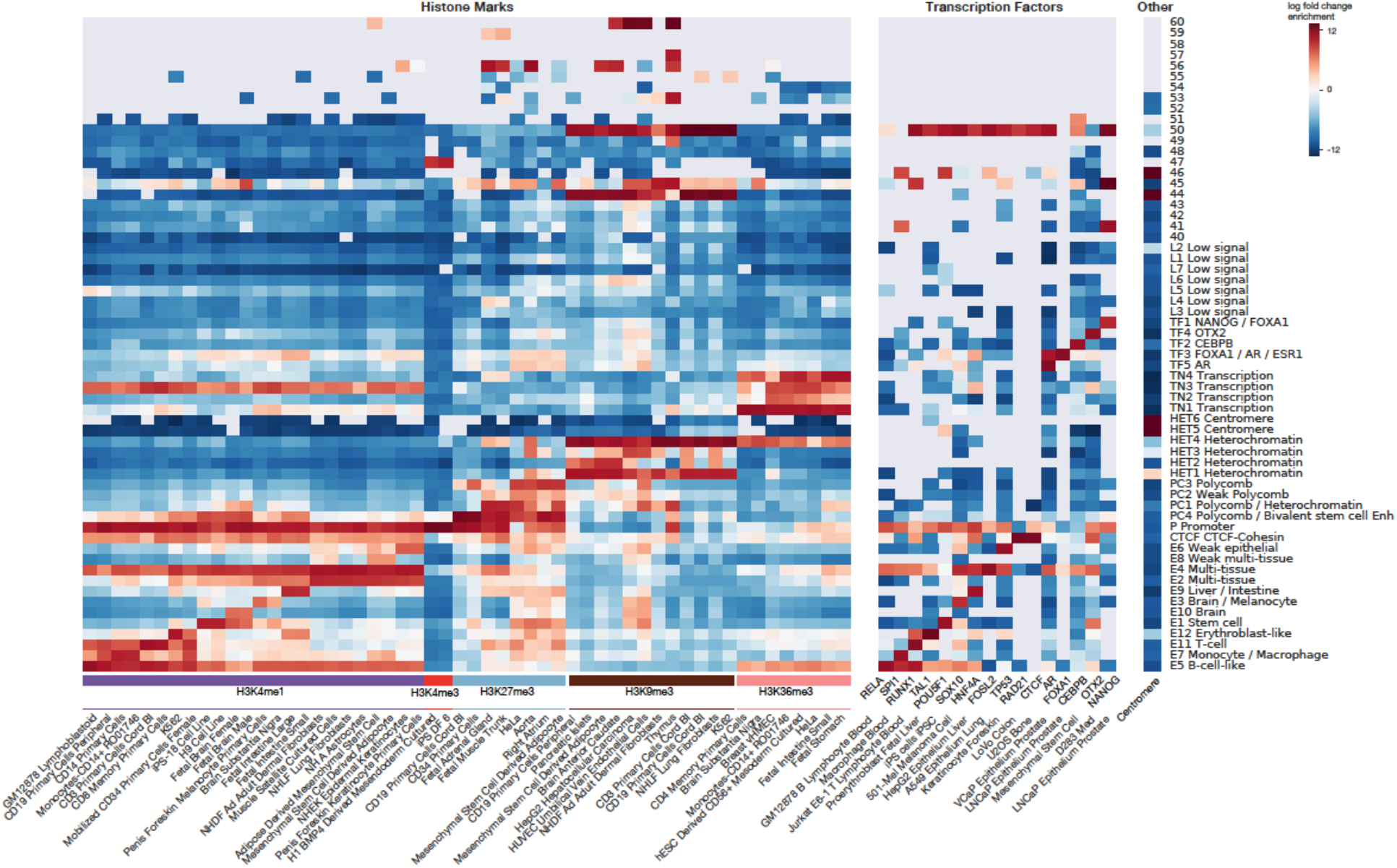
Enrichment of histone marks, transcription factors, and repeat annotations for the full set of 61 clusters output by Louvain community clustering. Log fold change enrichment over genome-average background is shown in the heatmap. No overlap is indicated by the gray color in the heatmap. Top 1-2 histone mark and TF annotation enrichments were selected for each sequence class.

**Supplementary File 1. The list of 21907 cis-regulatory profiles and Sei prediction performance.**

**Supplementary File 2. Summary of tissues and cell types covered by the Sei model.**

**Supplementary File 3. Top 25 enriched Cistrome Project chromatin profiles for each sequence class.** Log fold change enrichment over genome-average background is shown in the heatmap. No overlap is indicated by the gray color in the heatmap. We computed the enrichment for all 21,907 profiles predicted by Sei over 2 million random genomic positions. For each sequence class, the chromatin profiles are filtered to those having Benjamini-Hochberg corrected p-values (Fisher’s exact test, two-sided) < 2.2e-16 selecting the top 25 profiles based on log fold change enrichment.

**Supplementary File 4. Partitioned UKBB GWAS heritability by sequence classes using LDSR.** The proportions of heritability are represented by the LDSR estimate - 1 standard error, lower bounded by 0. The LDSR enrichment z-scores are also lower bounded by 0.

**Supplementary File 5. Significant UKBB GWAS trait - sequence class associations identified with LDSR conditioned on the baseline annotations.**

**Supplementary File 6. Predicted sequence class-level variant effects for HGMD regulatory disease mutations.** HGMD regulatory disease mutations with sequence-class level variant effect score >1.1 are included.

**Supplementary File 7. Detailed Sei model architecture specification.**

