## Supplementary figures and images for "A sequence-based global map of regulatory activity for deciphering human genetics"

### Supplementary Figure 1.pdf

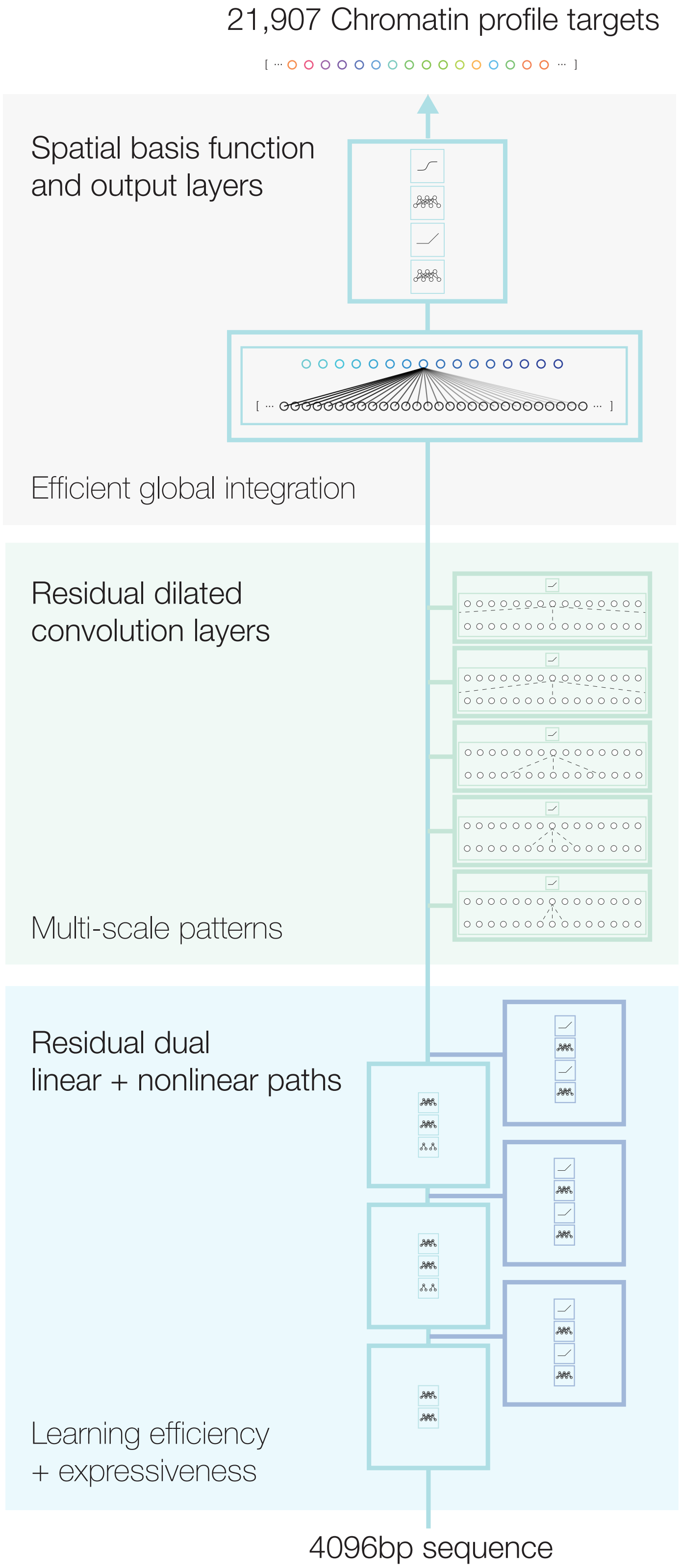

### Supplementary Figure 2.pdf

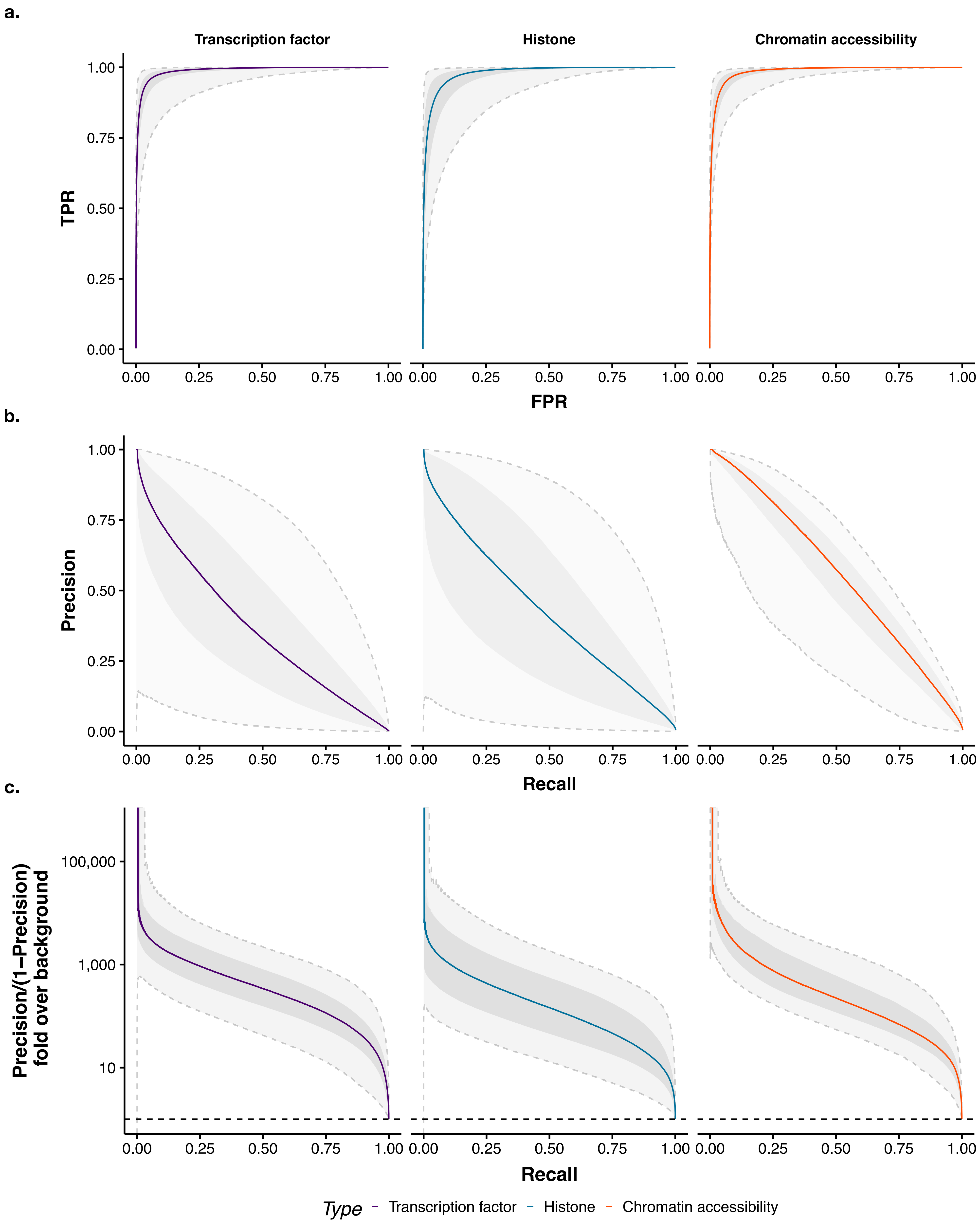

### Supplementary Figure 4.pdf

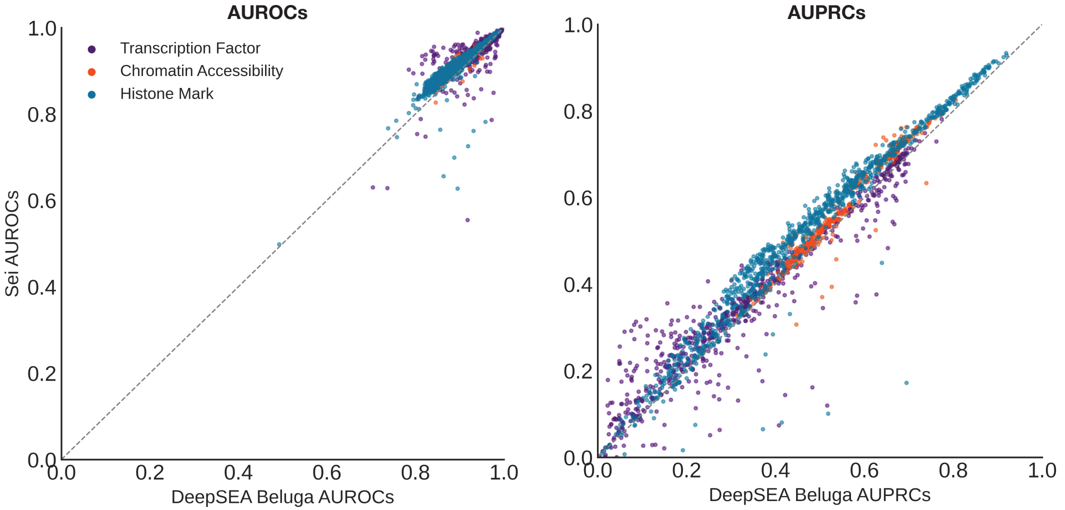

### Supplementary Figure 5.pdf

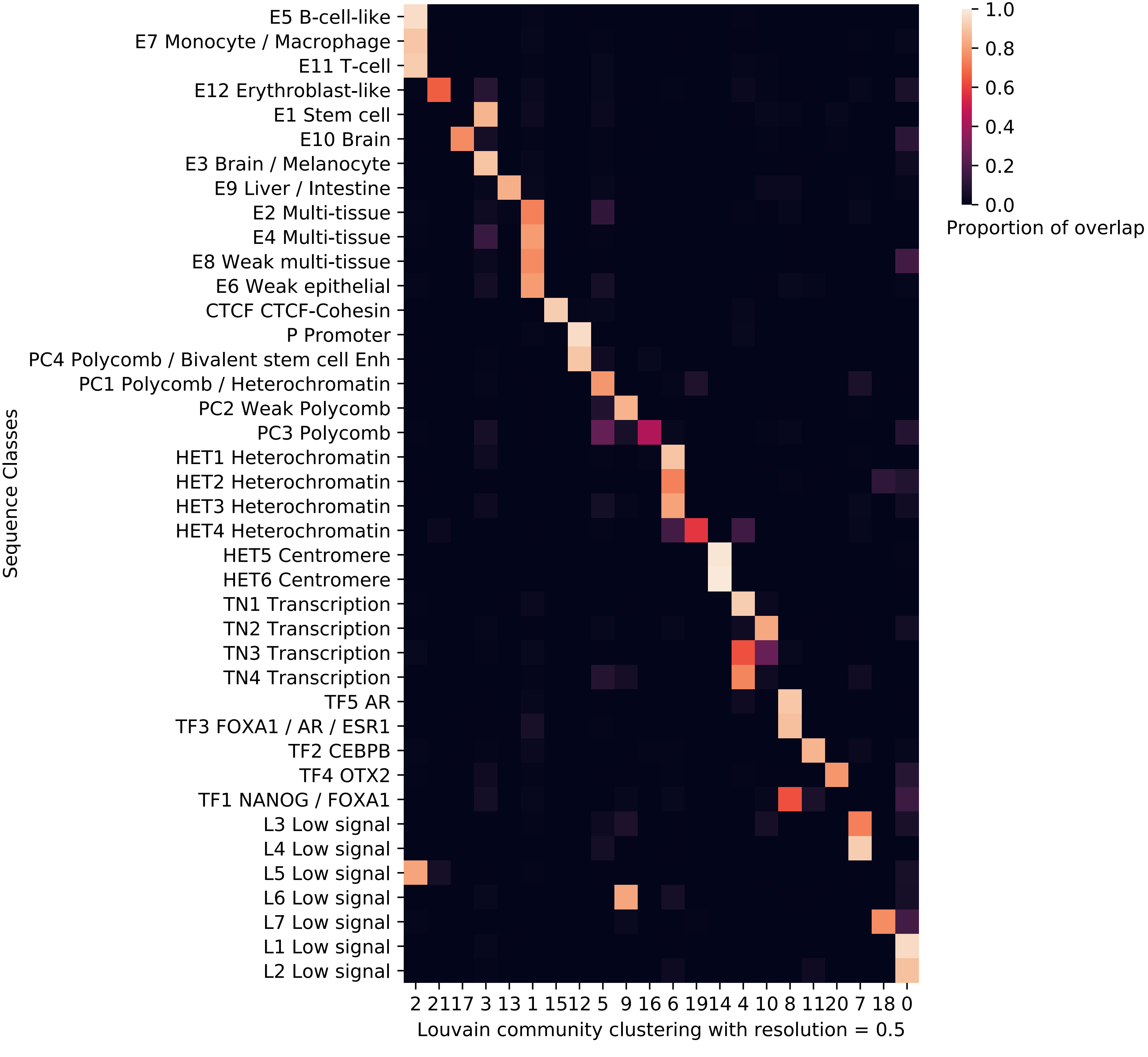

### Supplementary Figure 6.pdf

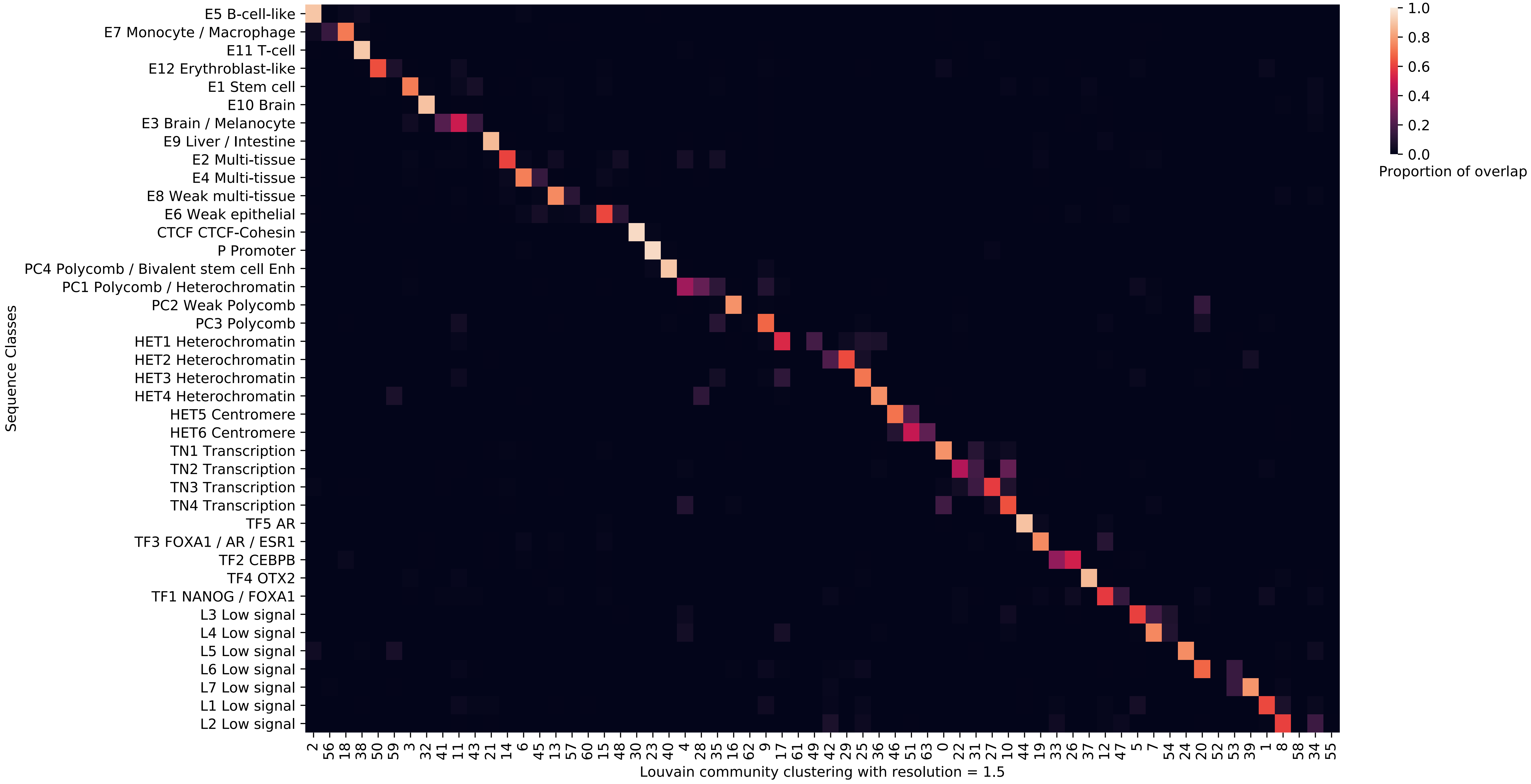

### Supplementary Figure 7 - H3K4me3.pdf

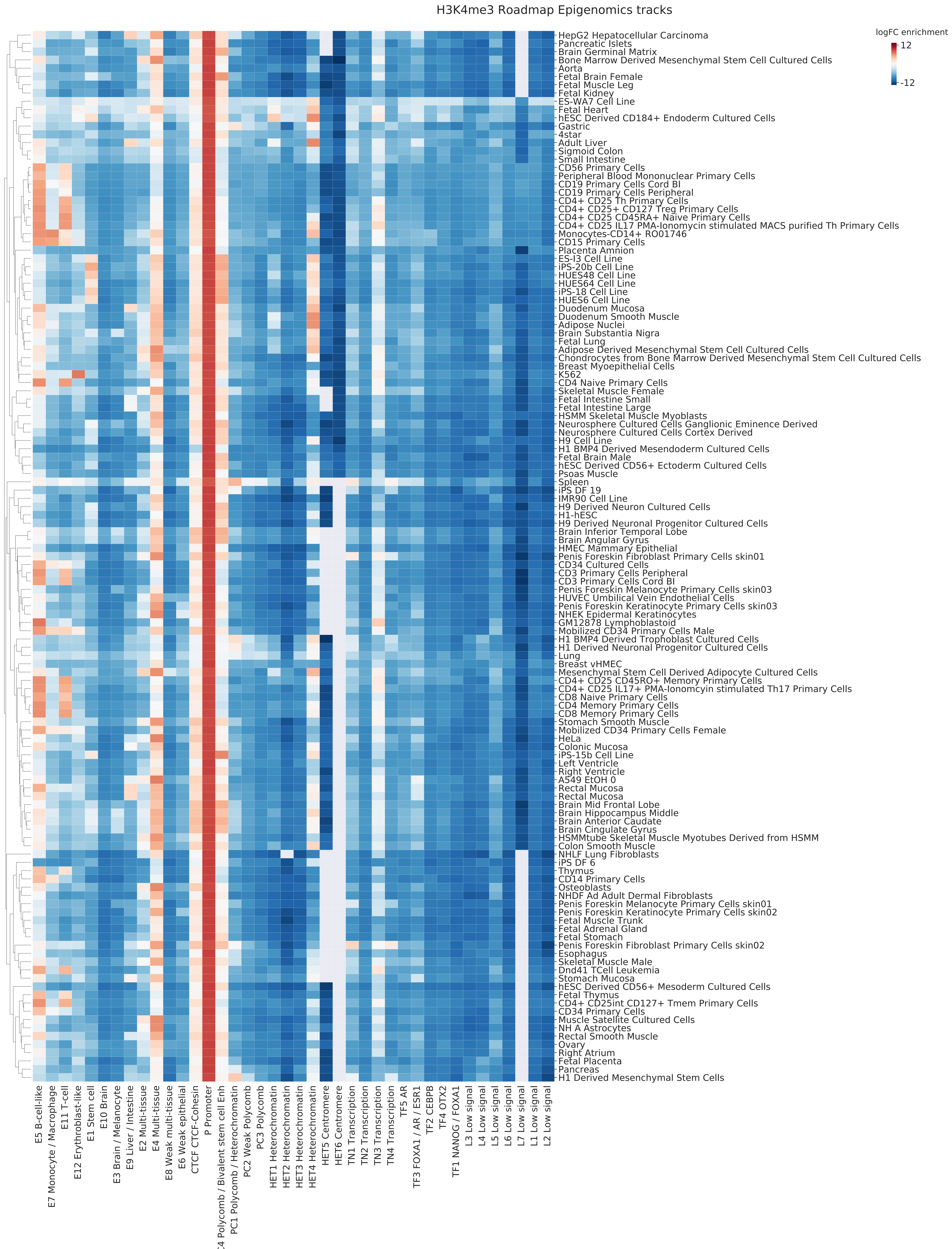

### Supplementary Figure 8 - H3K4me1.pdf

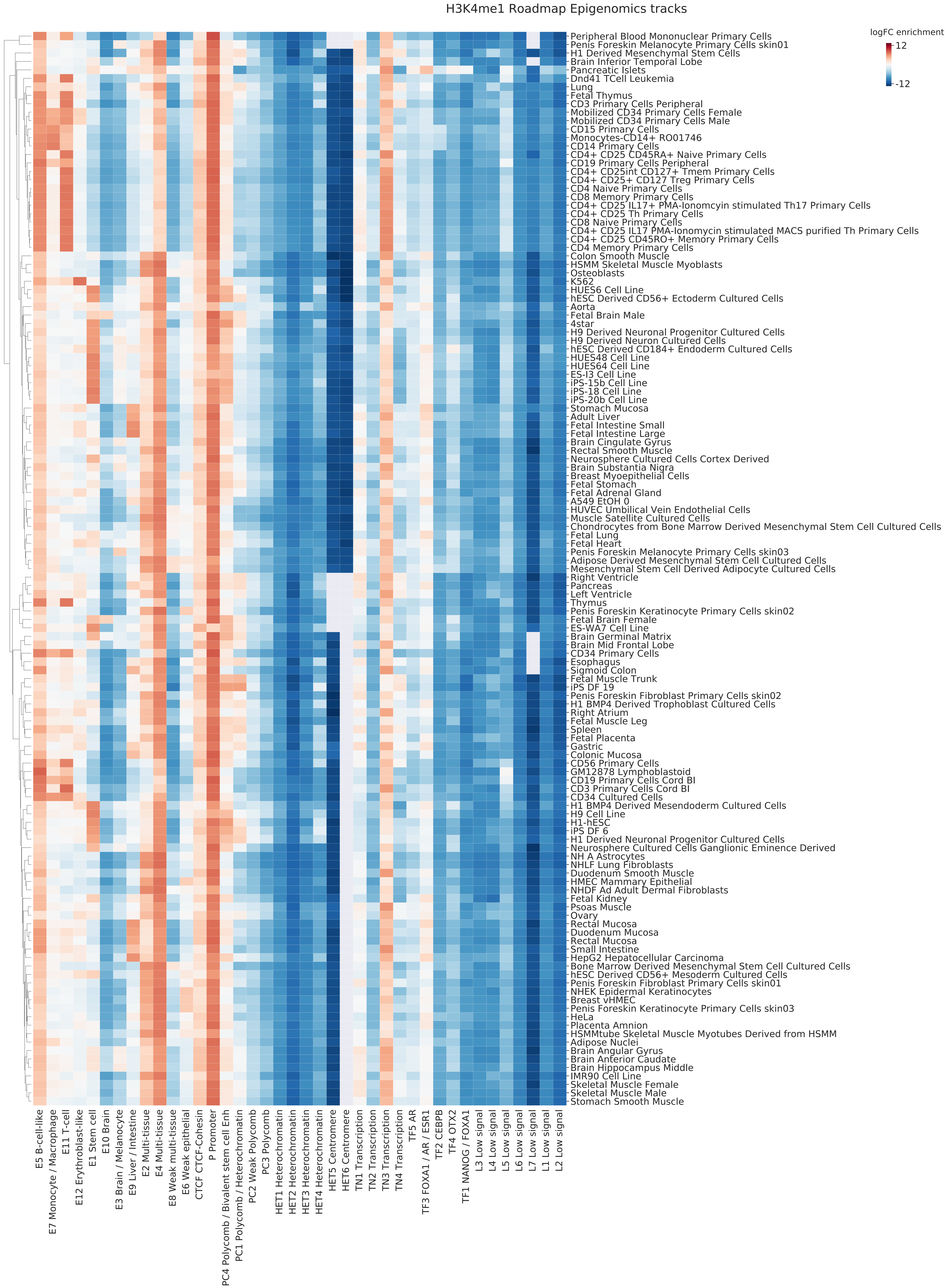

### Supplementary Figure 9 - H3K27ac.pdf

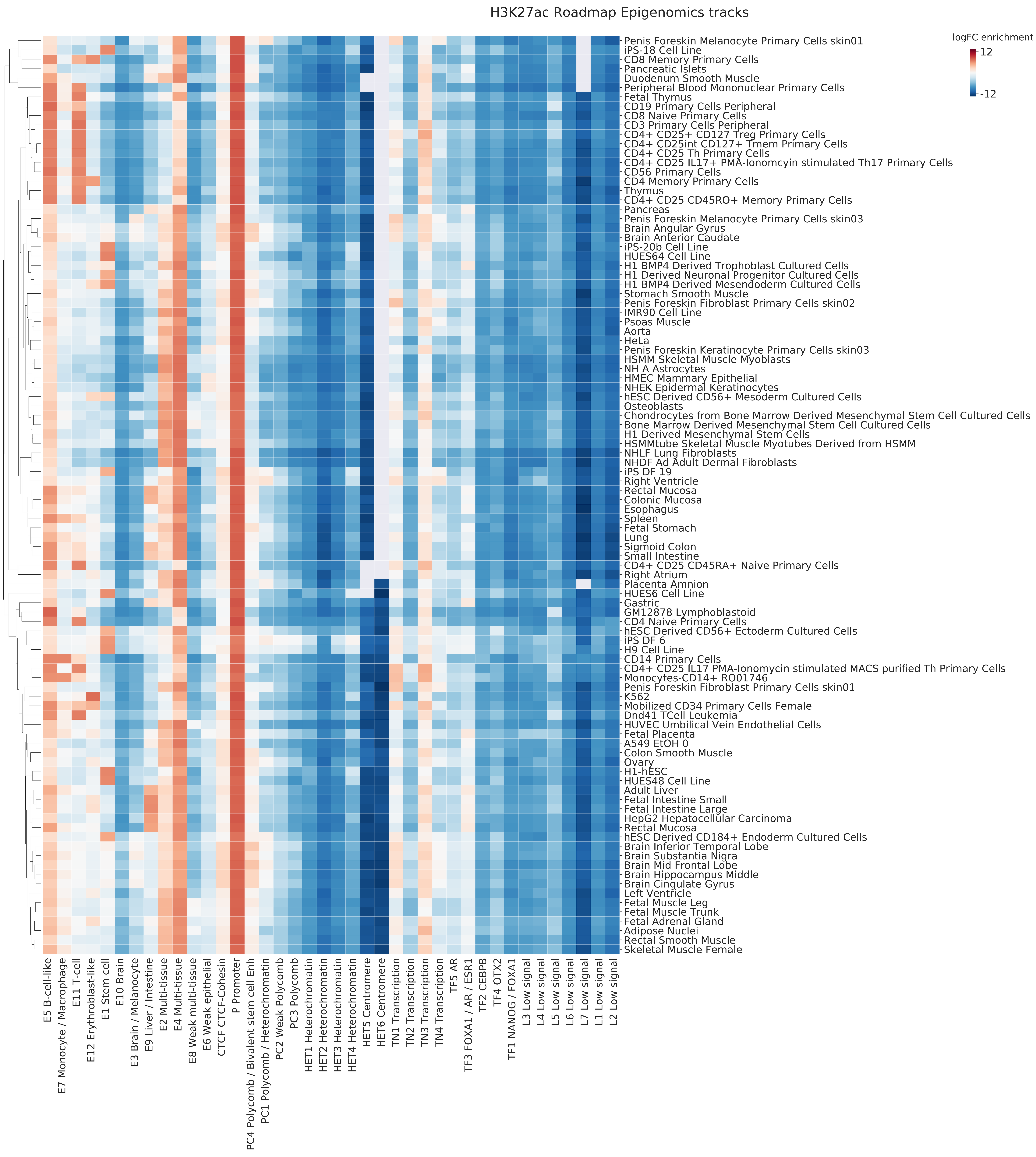

### Supplementary Figure 10 - H3K27me3.pdf

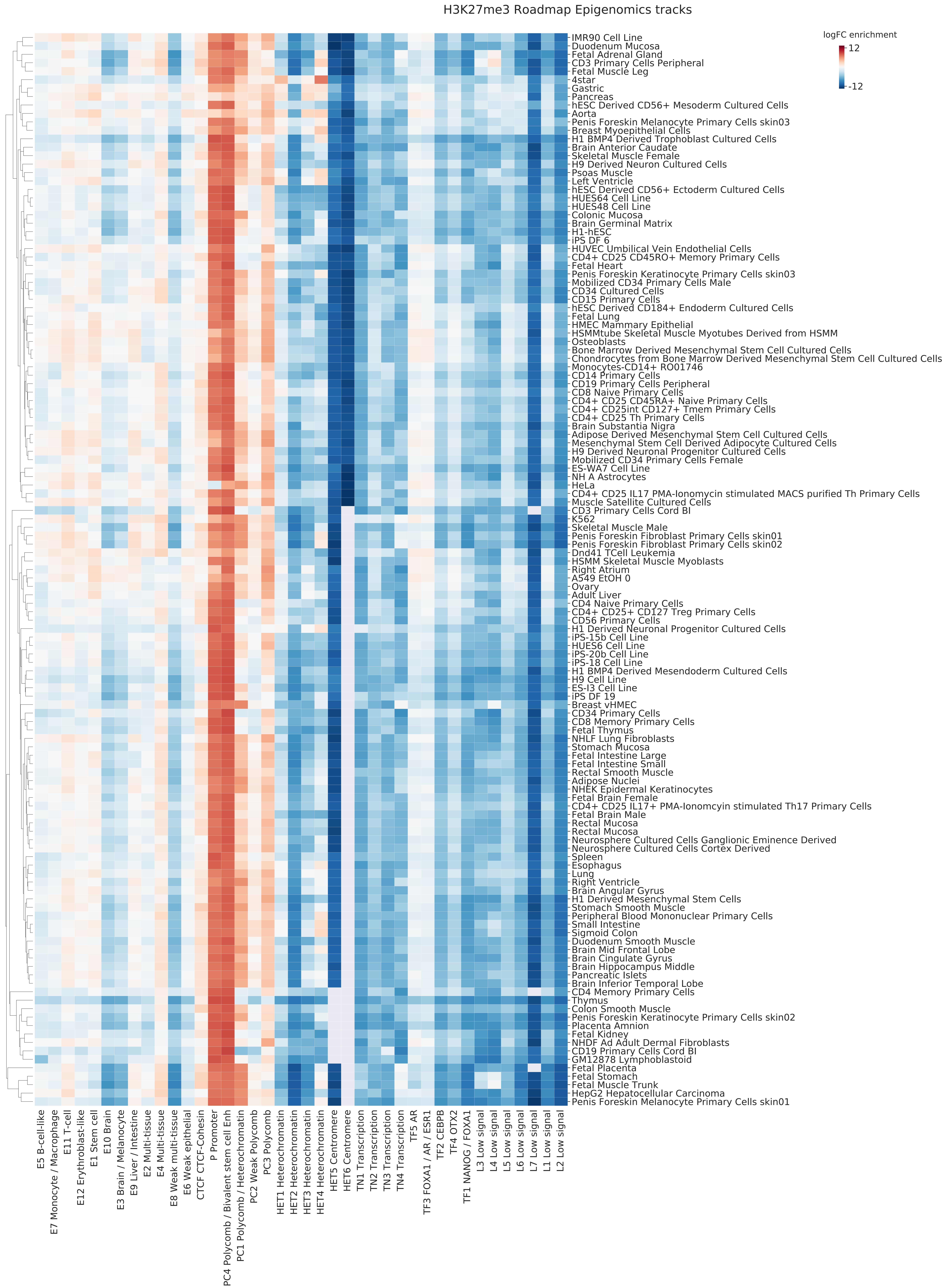

### Supplementary Figure 11 - H3K9me3.pdf

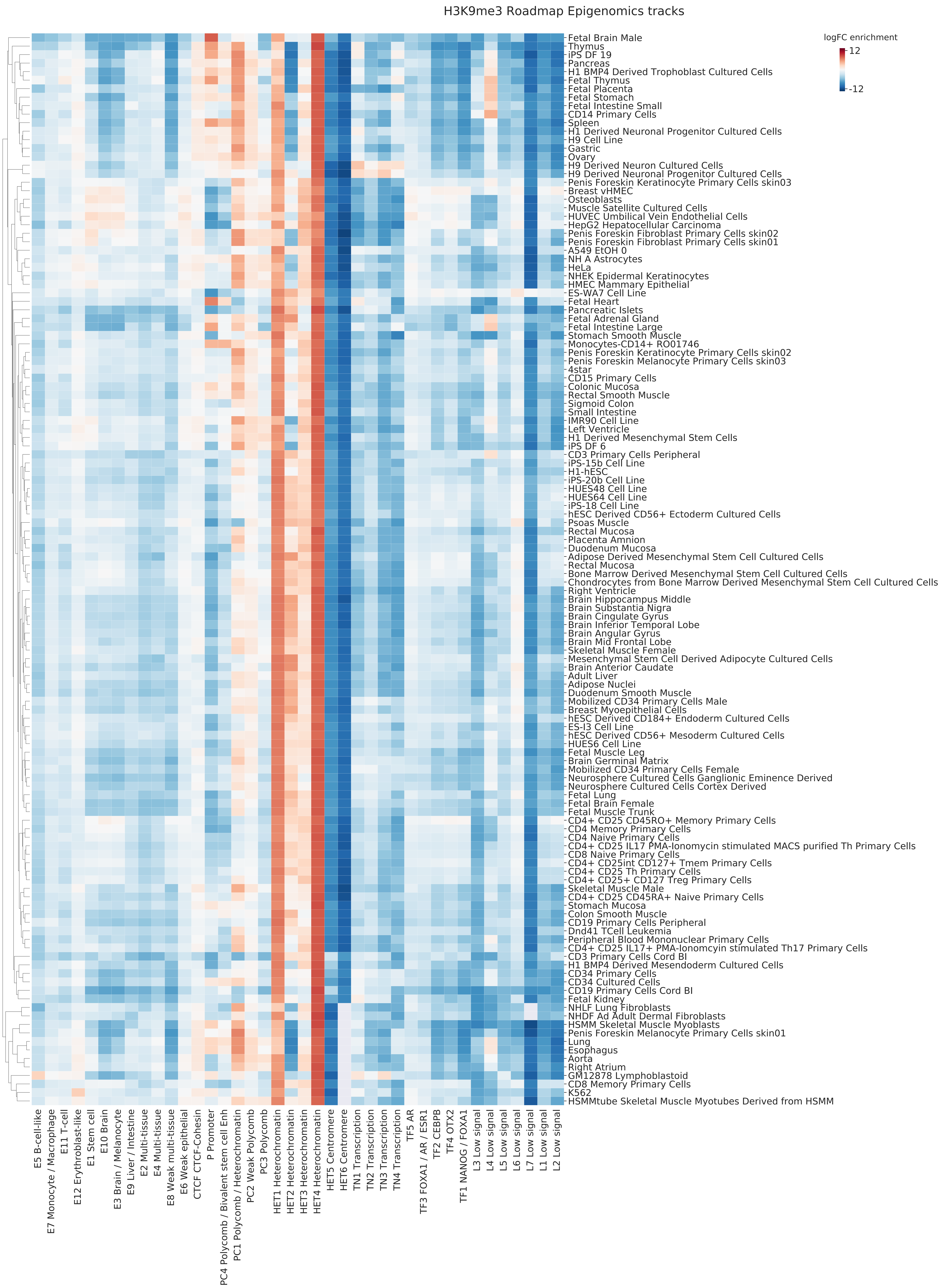

### Supplementary Figure 12 - H3K36me3.pdf

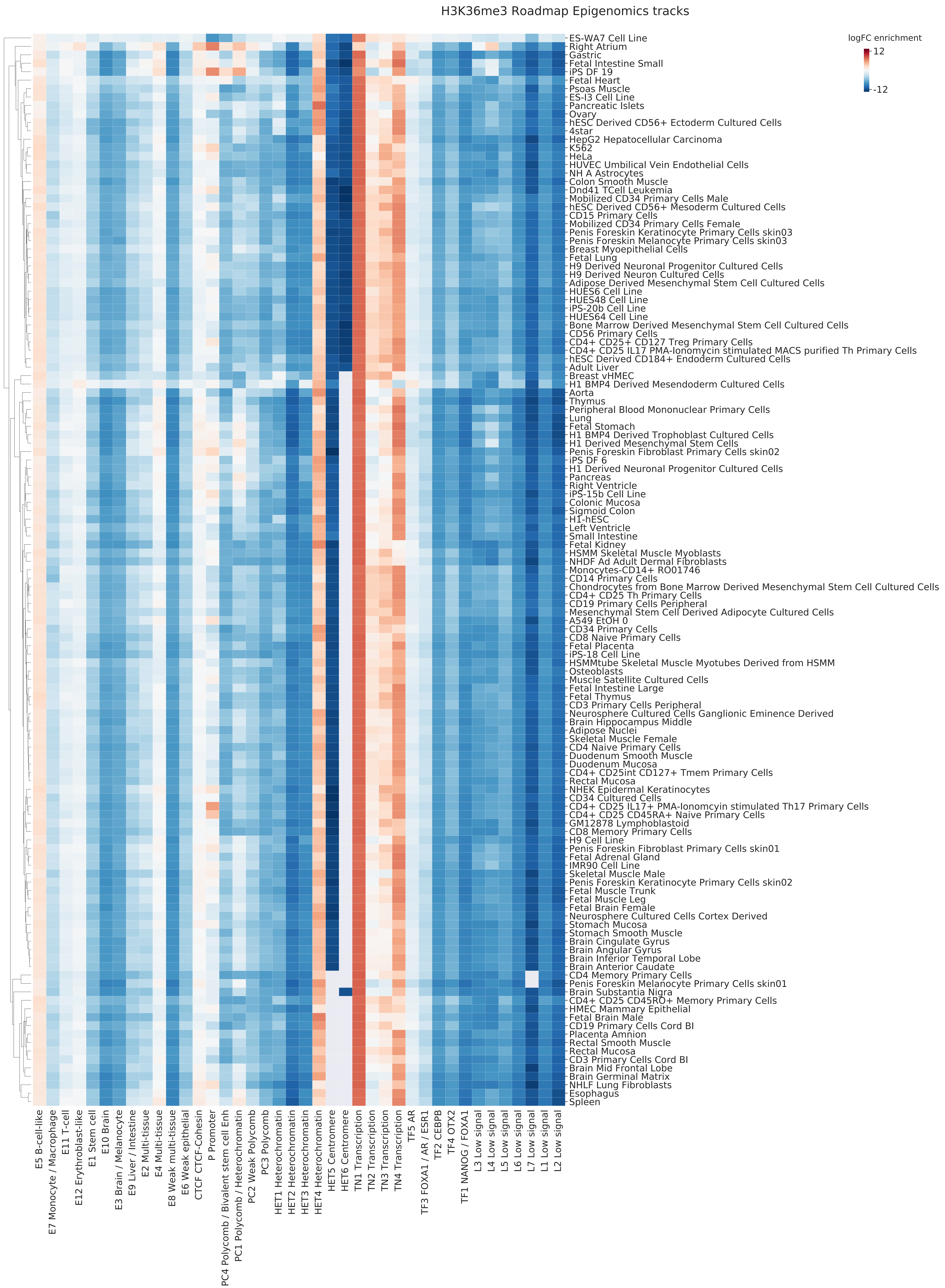

### Supplementary Figure 13.pdf

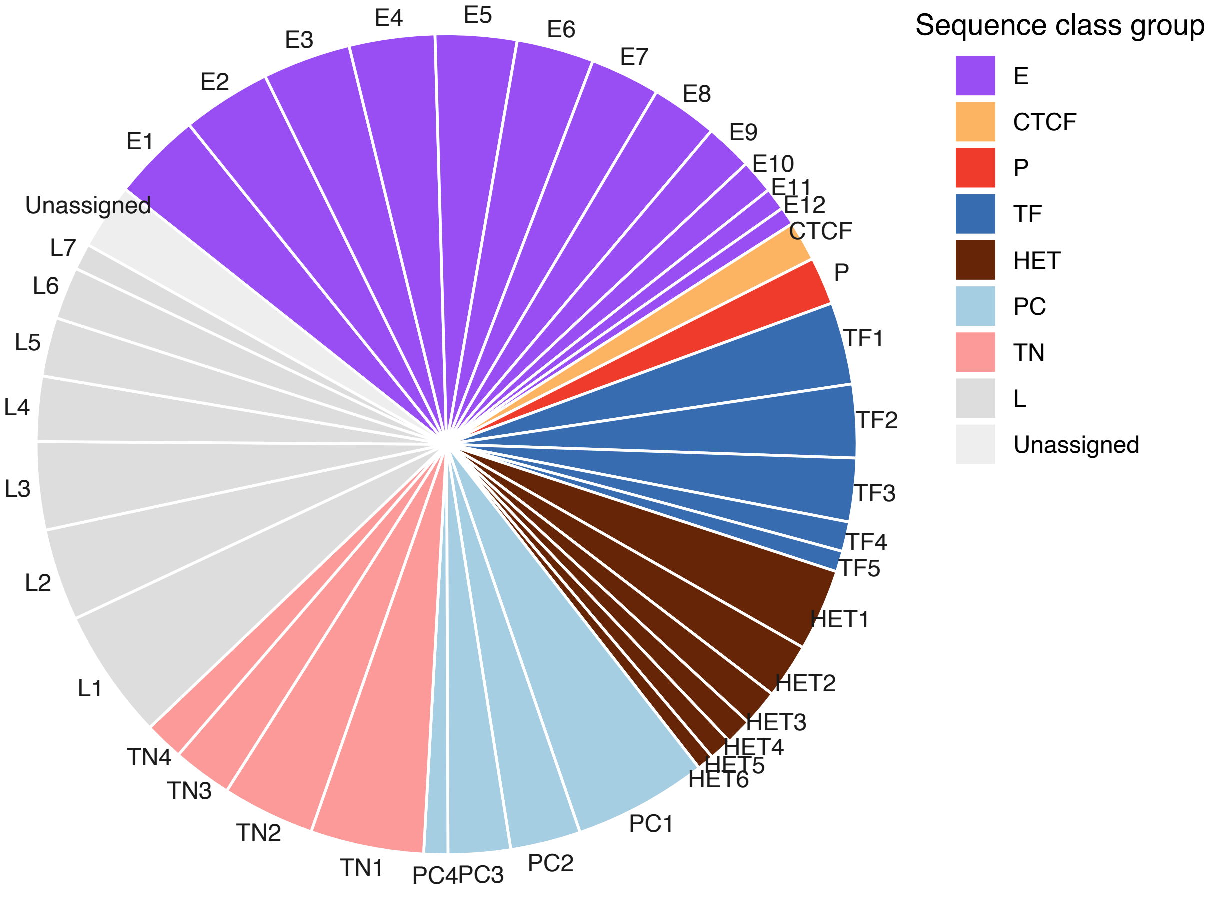

### Supplementary Figure 14.pdf

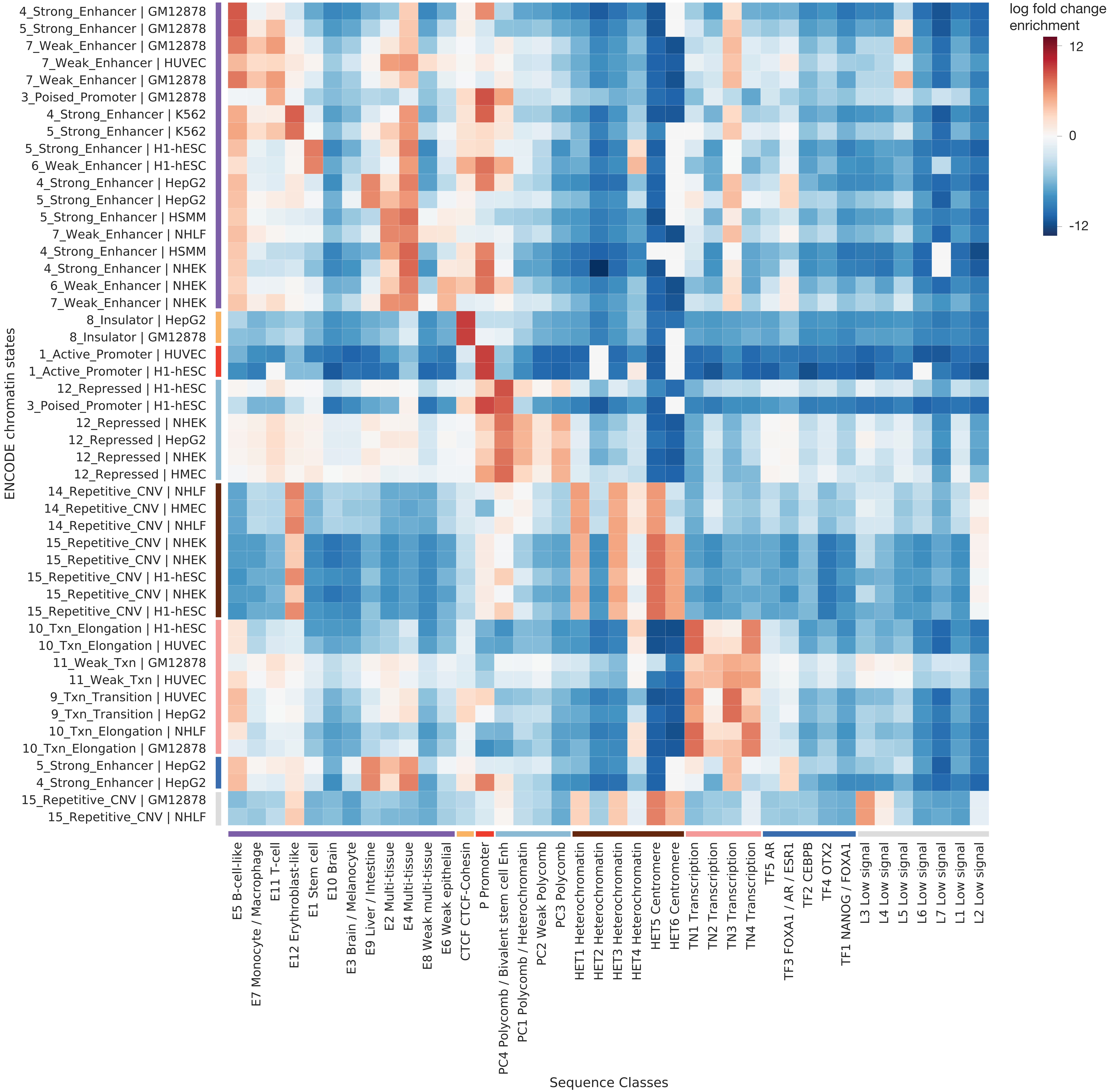

### Supplementary Figure 15.pdf

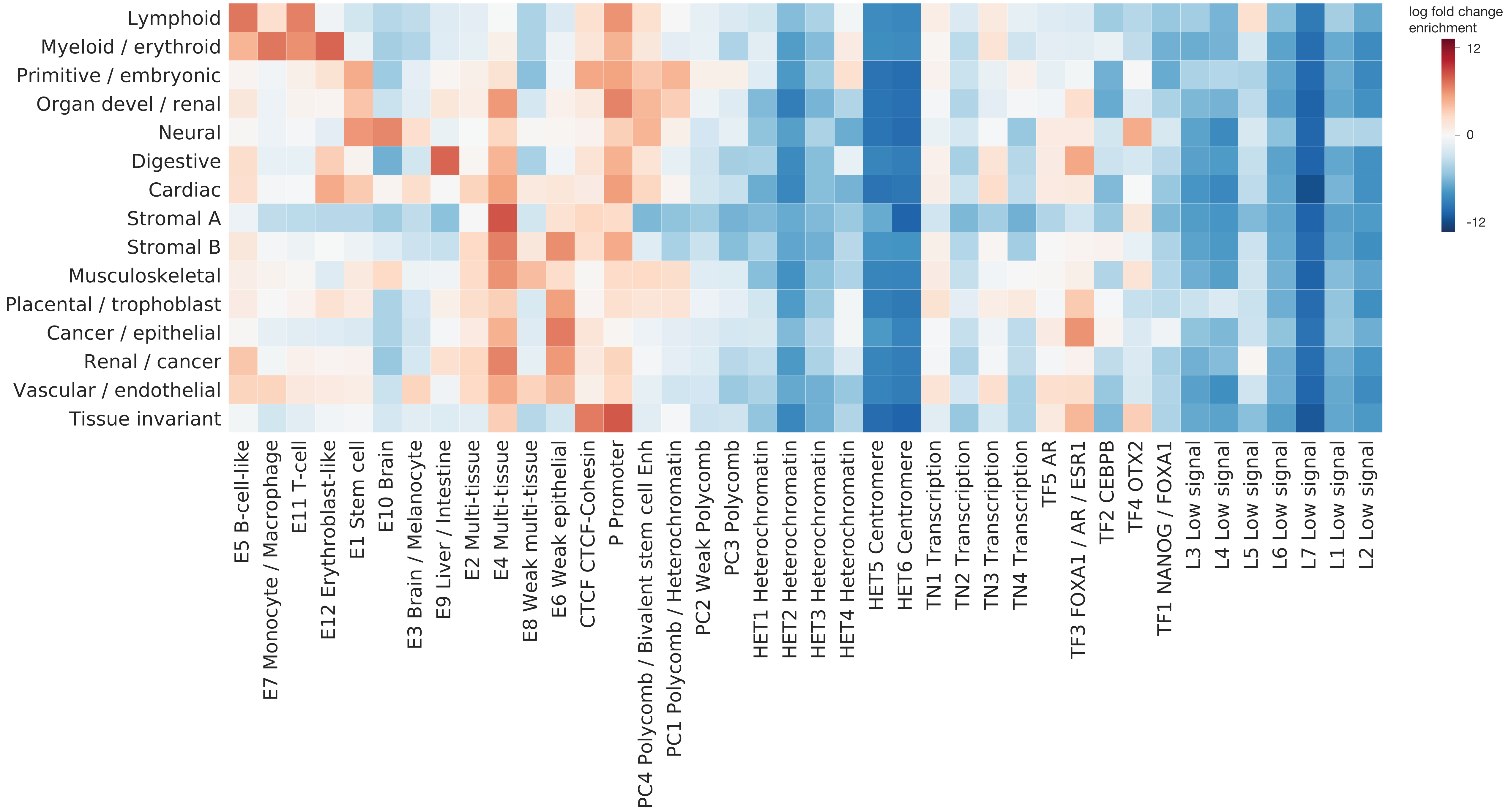

### Supplementary Figure 16.pdf

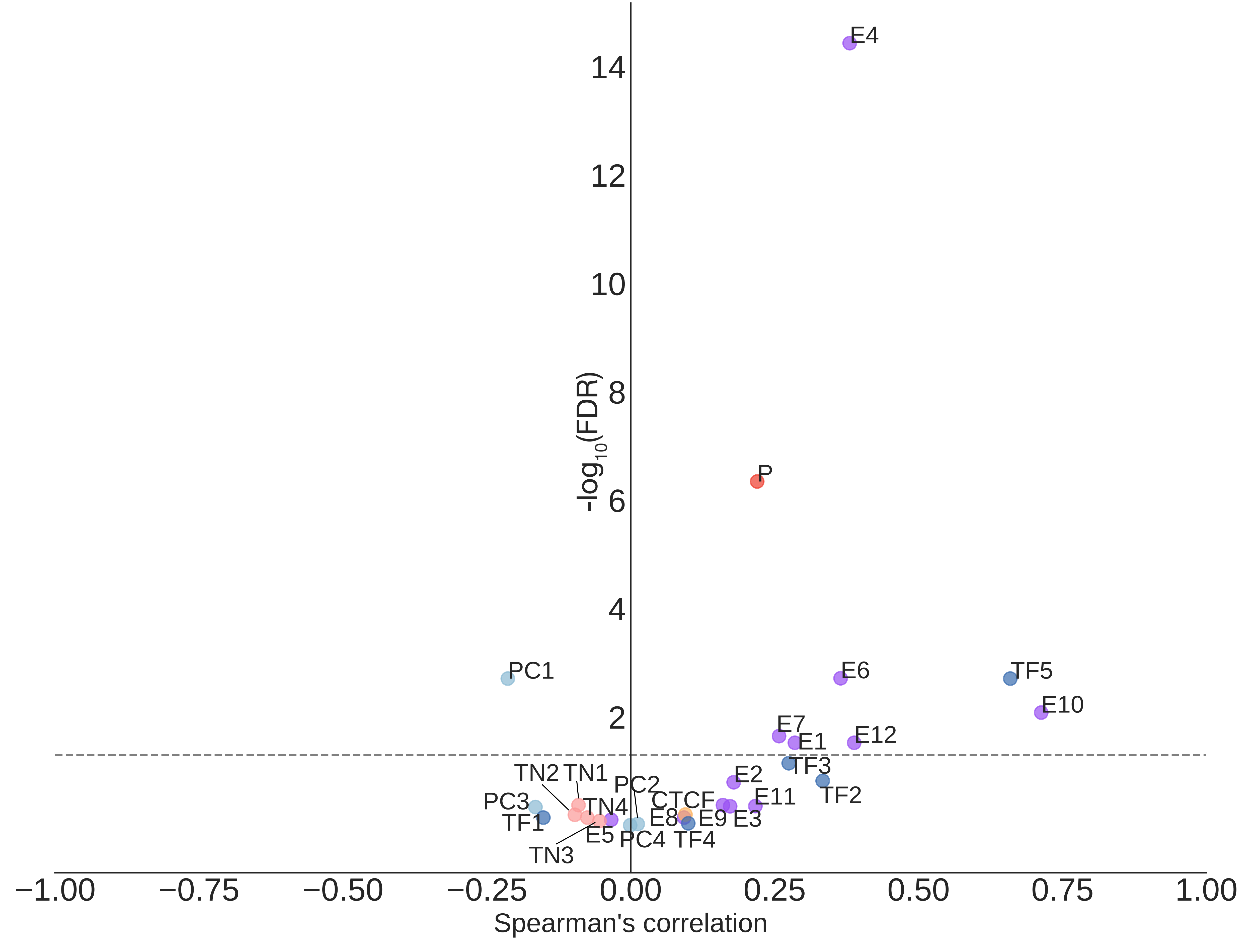

### Supplementary Figure 17.pdf

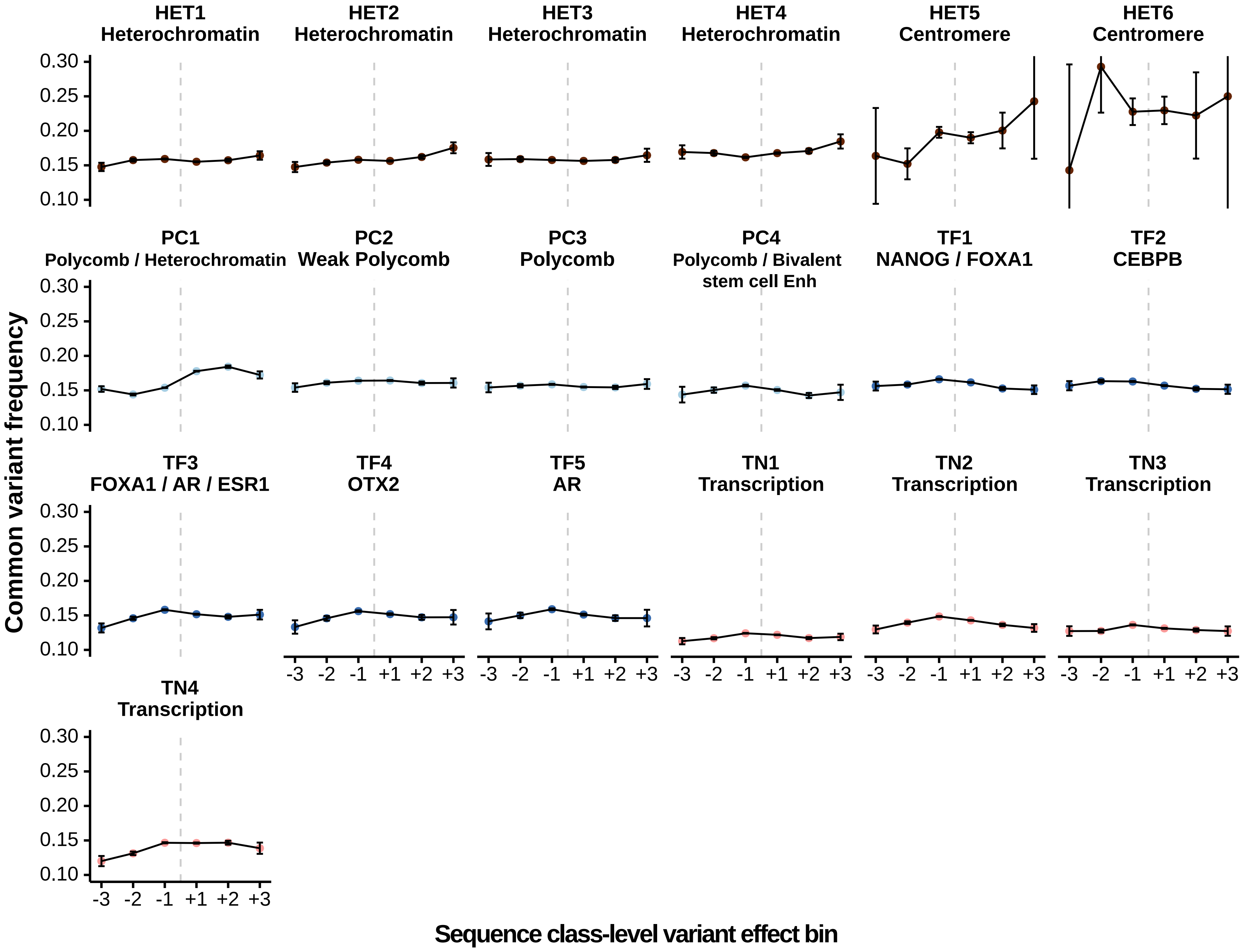

### Supplementary Figure 18.pdf

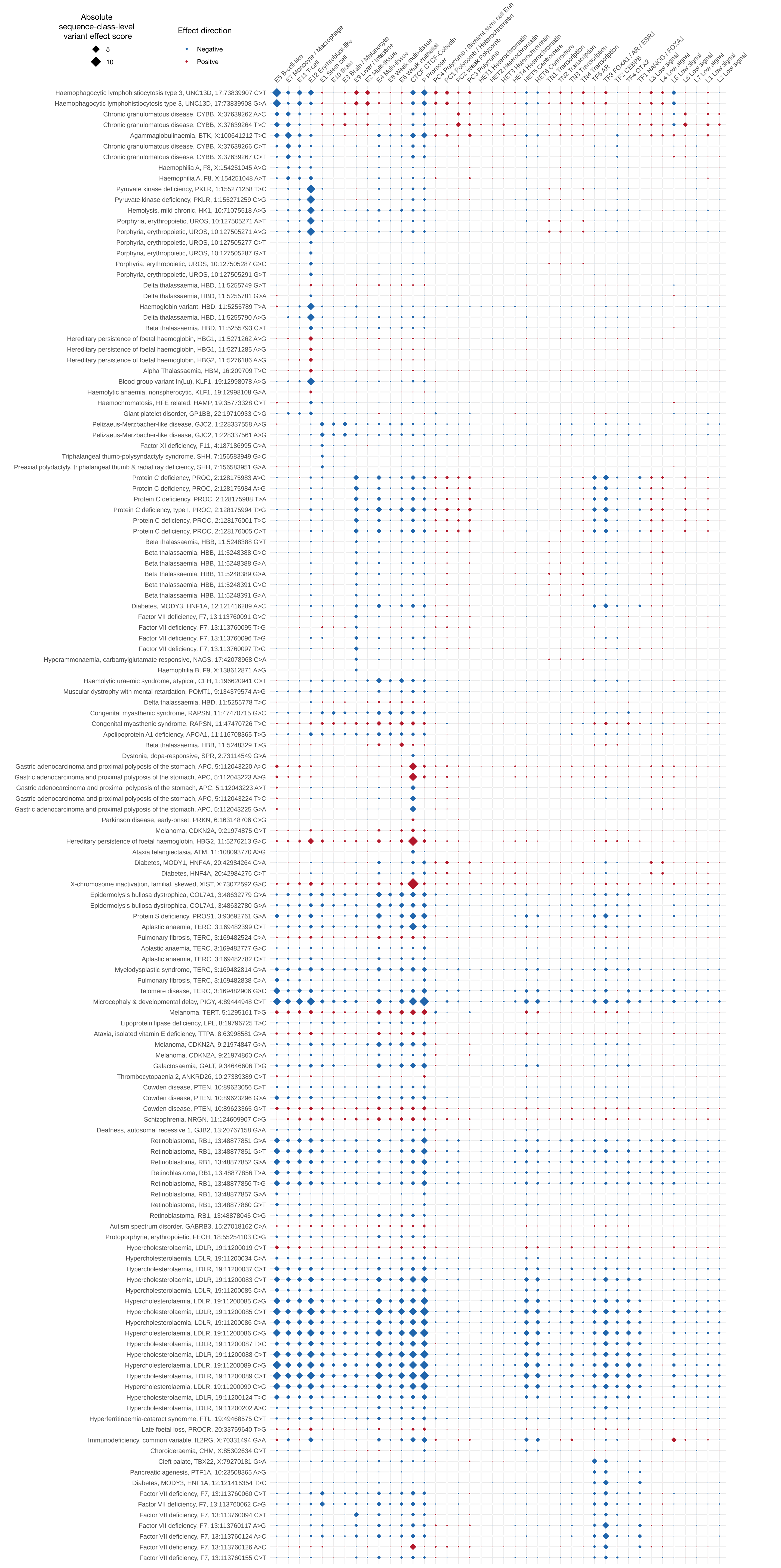

### Supplementary Figure 19.pdf

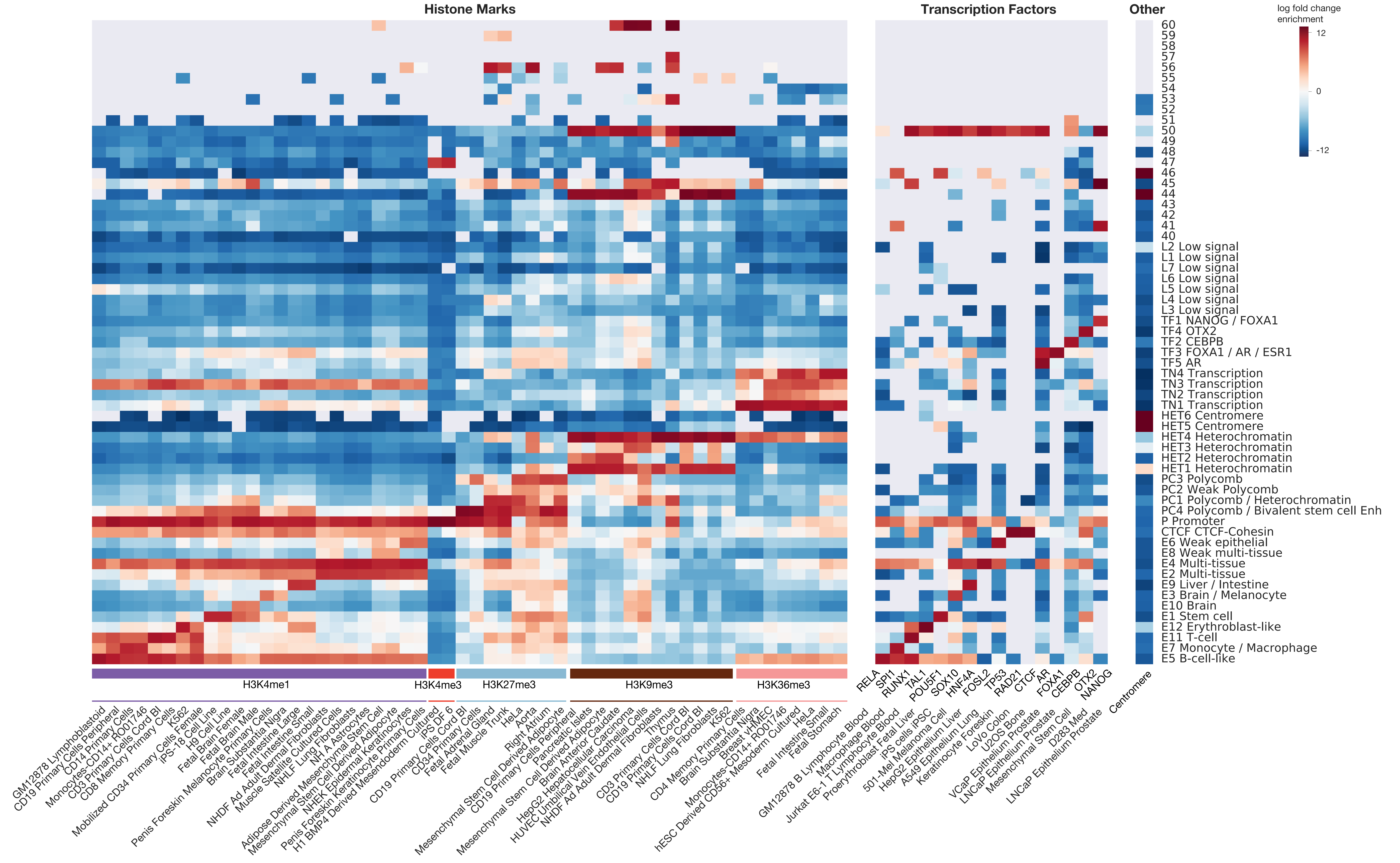

### Supplementary File 3.pdf

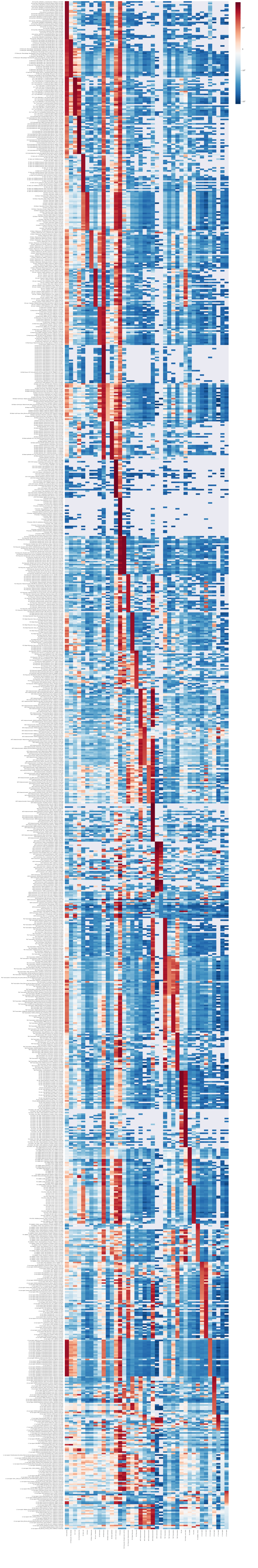

### Supplementary File 7.pdf

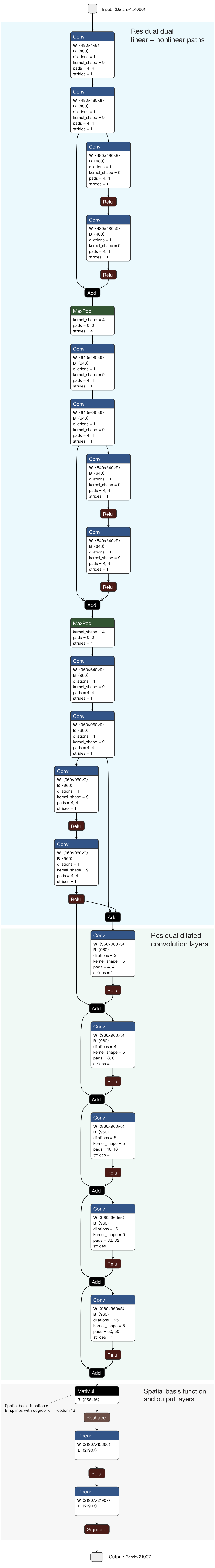
